# Regeneration of distinct complex structures in the annelid *Platynereis* is partially based on common morphological, cellular, and molecular events

**DOI:** 10.1101/2025.10.03.680064

**Authors:** Zoé Velasquillo Ramirez, Viktor Starunov, Louis Paré, Loeiza Baduel, Théaud Hezez, Sophie Lemoine, Catherine Senamaud-Beauford, Michel Vervoort, Morgane Thomas-Chollier, Nikos Konstantinides, Eve Gazave

## Abstract

Many metazoan species harbor fascinating regenerative capabilities, but the underlying mechanisms remain poorly understood. Whether the capacity to regenerate complex structures successfully relies on common or divergent features is still an open question. To identify the key molecular and cellular elements necessary for successful regeneration, we investigated the efficient regeneration of two distinct complex structures in the annelid worm *Platynereis dumerilli*. By combining classical morphological and developmental approaches, with state-of-the-art single-cell RNA sequencing and analysis, we conducted a comprehensive comparison of locomotory appendage and posterior part regeneration. We uncovered the rich cell type diversity of this *Platynereis* appendage, over a third of which are nerve-related populations, highlighting the importance of its sensory function. We also described its regenerative process at the morphological and cellular levels, defining stages that include the formation of a proliferative blastema. We then compared posterior part and appendage blastemas, specifically assessing their cell diversity and cell differentiation trajectories. We found transcriptionally similar epithelial and mesodermal progenitors at play during posterior and appendage regeneration, although their specific trajectories differed to some extent. Our work, revealing partial morphological, molecular and cellular parallels between these two efficient regenerations within a single species, sets the foundation for addressing the fundamental question of regeneration success in animals.

## Introduction

Restorative regeneration, the ability to reform a lost or damaged body part is a phenomenon found in members of all major lineages of Metazoa (Bely & Nyberg, 2010; Bideau et al., 2021; Poss, 2010). The potential for regeneration varies greatly among different species, ranging from the regeneration of scarce cell types to the renewal of organs, complex structures, and even the entire body (Bideau et al., 2021; Grillo et al., 2016). During successful regeneration events, the animals restore their body plan, remake many different cell types/tissues, synchronize their dynamics and fate to integrate them with those that remained, and rebuild a functional structure (Poss & Tanaka, 2024). Why and how some species can successfully regenerate complex parts of their body, while others cannot, is still a major open question in the field (Gazave & Röttinger, 2021; Poss & Tanaka, 2024). Expanding the number of regeneration model organisms by studying new species is advocated as indispensable to solving this long-standing question (Sasidharan & Sánchez Alvarado, 2022). Hence, cross-species analysis of regeneration among metazoans have revealed recurrent players at the genetic, epigenetic and post-transcriptional levels (Fumagalli et al., 2018; Khyeam et al., 2021; Rouhana & Tasaki, 2016; Vullien et al., 2025). However, finding commonalities for successful regeneration between different species remains a major challenge. Indeed, the regeneration processes rarely show a comparable timing and staging system and the structures considered *de facto* differ a lot, as does the genetic complexity of the species compared (Srivastava, 2021). One powerful means to reduce the complexity of this question is to focus on a single species that can regenerate different complex structures, to assess and compare the morphological, cellular, and molecular actors involved. Hence, one can determine if successful regenerations of different structures rely on similar or divergent processes.

Annelids, or segmented worms, represent a major group of highly regenerative species, surprisingly understudied in modern regenerative biology (Kostyuchenko & Kozin, 2021; Özpolat & Bely, 2016). Known to regenerate their very complex posterior and/or anterior ends, appendages, and a diversity of sensory structures, they represent an ideal biological playground for understanding the success of complex structure regeneration (Hyman, 1940; Özpolat & Bely, 2016). Among annelids, one key model species, *Platynereis dumerilii* has (re)emerged as a powerful model due to its extended regenerative capabilities (Hofmann, 1976; Özpolat et al., 2021; Schenkelaars & Gazave, 2021; Vervoort & Gazave, 2022). Upon amputation of the posterior part of their body, removing the terminal part of the worm (named pygidium), the stem cell-rich subterminal growth zone (GZ, responsible for the continuous growth of the worms (Gazave et al., 2013)) and several segments, *Platynereis* worms regenerate those missing structures in five days through posterior regeneration (Planques et al., 2019). Similar to most regeneration processes, *Platynereis’* posterior regeneration can be divided into three sequential steps (Bideau et al., 2021; Tiozzo & Copley, 2015). First, the wound heals through the formation of a wound epithelium, within a day. The second step relies on a regeneration-specific structure, called a blastema. The blastema is formed within 2 to 3 days post amputation (dpa), thanks to the mobilization of precursor cells of local origin which are most likely produced through dedifferentiation (Bideau et al., 2024; Stockinger et al., 2024). The last morphogenetic step (3 to 5 dpa) consists not only in the reestablishment of the differentiated tissues of the pygidium but also of the GZ stem cells (Bideau et al., 2025), which in turn will produce new segments (Planques et al., 2019). In addition, *Platynereis* is able to regenerate another important body structure: its locomotory, respiratory, and sensory body paired appendages named parapodia (Grimmel et al., 2016; Paululat & Purschke, 2025). Whether parapodial and posterior regenerations in *Platynereis* rely on similar or different processes remains an open question.

Here, we sought to identify the molecular and cellular programs underpinning the regeneration success of complex structures in *Platynereis.* To this end, we made a thorough comparison between parapodial and posterior regeneration combining classical morphological and developmental approaches, with state-of-the-art single-cell RNA sequencing and analysis. Firstly, we profiled the adult cell populations of parapodia, uncovering the extent of their diversity, and experimentally validated them by studying their expression patterns. Then, we conducted a detailed characterization of parapodial regeneration. We defined morphological stages, studied the cell proliferation dynamics, and the kinetics of the reformation of the various tissues and structures of the parapodia. We also generated a transcriptomic cell atlas for the key parapodial regeneration stage: the blastema. We identified many progenitor populations and reconstructed their differentiation trajectories. Finally, we capitalized on the integration of newly produced and already available atlases for posterior regeneration blastema (Stockinger et al., 2024), as well as important knowledge gathered over the years on this process (Bideau et al., 2024, 2025; Novikova et al., 2025; Paré et al., 2023; Pfeifer et al., 2012; Planques et al., 2019, 2021; Vervoort & Gazave, 2022), to compare cell type diversity and differentiation trajectories between parapodial and posterior regeneration. We provide evidence for the use of transcriptionally similar epithelial and mesodermal progenitors during posterior and parapodial regeneration. However, their associated differentiation trajectories, although restricted to the blastema stage, are partially different. Hence, our study comparing different types of regeneration within a unique species, allowing the identification of common elements at play, set the foundation for addressing the fundamental question of regeneration success in animals.

## Results

### *Platynereis* juvenile parapodia featured at an unprecedented resolution: from morphology to cell-type atlas

In the juvenile *Platynereis*, parapodia are proximo-distal bifurcated paired appendages flanking the segments of the trunk region, as is typical of annelids (Fig. 1A). Biramous, they are composed of a dorsal notopodium and a ventral neuropodium. Each ramus is composed of a cirrus, a lobe and a beam of extracellular chitineous structures named chaetae (Fig. 1B_1_, B_5_) (Grimmel et al., 2016; Schenkelaars & Gazave, 2021; Verdonschot, 2015). Histological sections confirmed a complex internal structure with two acicula, a robust skeletal chaeta stabilizing each lobe, and several types of glands. In the first ten segments, parapodia bear spinning glands secreting materials for worm cocoon synthesis (Fischer et al., 2010). In more posterior segments, notopodia bear flask-shaped glands and both lobes present elongated tubular-cell glands on the tips of their lobes (Daly, 1973) (Fig. 1B_2_, dark blue; Supp. Fig. 1A_1_-A_3_). Like many other annelids, *Platynereis* also presents a pair of excretory organs per segment (Fig. 1B_2_, orange). These organs, named metanephridia, pick up bodily fluids through a ciliated funnel and reabsorb reclaimable components before releasing waste through a pore at the base of the neuropodia (Hasse et al., 2010). As a major sensory structure, parapodia are highly innervated with a parapodial ganglia present at the base of the appendages from which three main nerves originate (Fig. 1B_3_, B_5_, green, Supp. Fig. 1B). The first bifurcates at the ventral side of the parapodia to innervate both the cirri and the tubular-cell glands, as well as the muscles at the base of the ventral chaeta and acicula. The second innervates the dorsal lobe. The third branches out to innervate the dorsal cirri, the tubular-cell and flask-shaped glands, as well as the base of the chaetal and acicular muscles. This third nerve then loops to innervate the dorsal gland found at the base of the parapodia. In line with their locomotive function, parapodia also have a dense network of muscles found in the two lobes (Fig. 1B_4_, B_5_, pink, Supp. Fig. 1C_1_-C_3_). Among the major recognizable muscles, acicular muscles connect the base of the acicula respectively to the ventral and dorsal body walls (Supp. Fig. 1C_1_, orange), while inter-acicular muscles connect the base of both acicula (Supp. Fig. 1C_1_, yellow). The acicula are also attached to the chaetal sacs through chaetal sac retractor muscles (Supp. Fig. 1C_1_, dark blue). Additional muscles connect the inter-acicular muscles to both lobes (Supp. Fig. 1C_2_, red). Chaetal protractor muscles link the base of the chaetal sacs to the tips of their respective lobes (Supp. Fig. 1C_2_, light blue). Finally, ventral and dorsal muscles connect to both the ventral and dorsal lobes (Supp. Fig.1C_3_, pink and purple).

**Figure 1.**
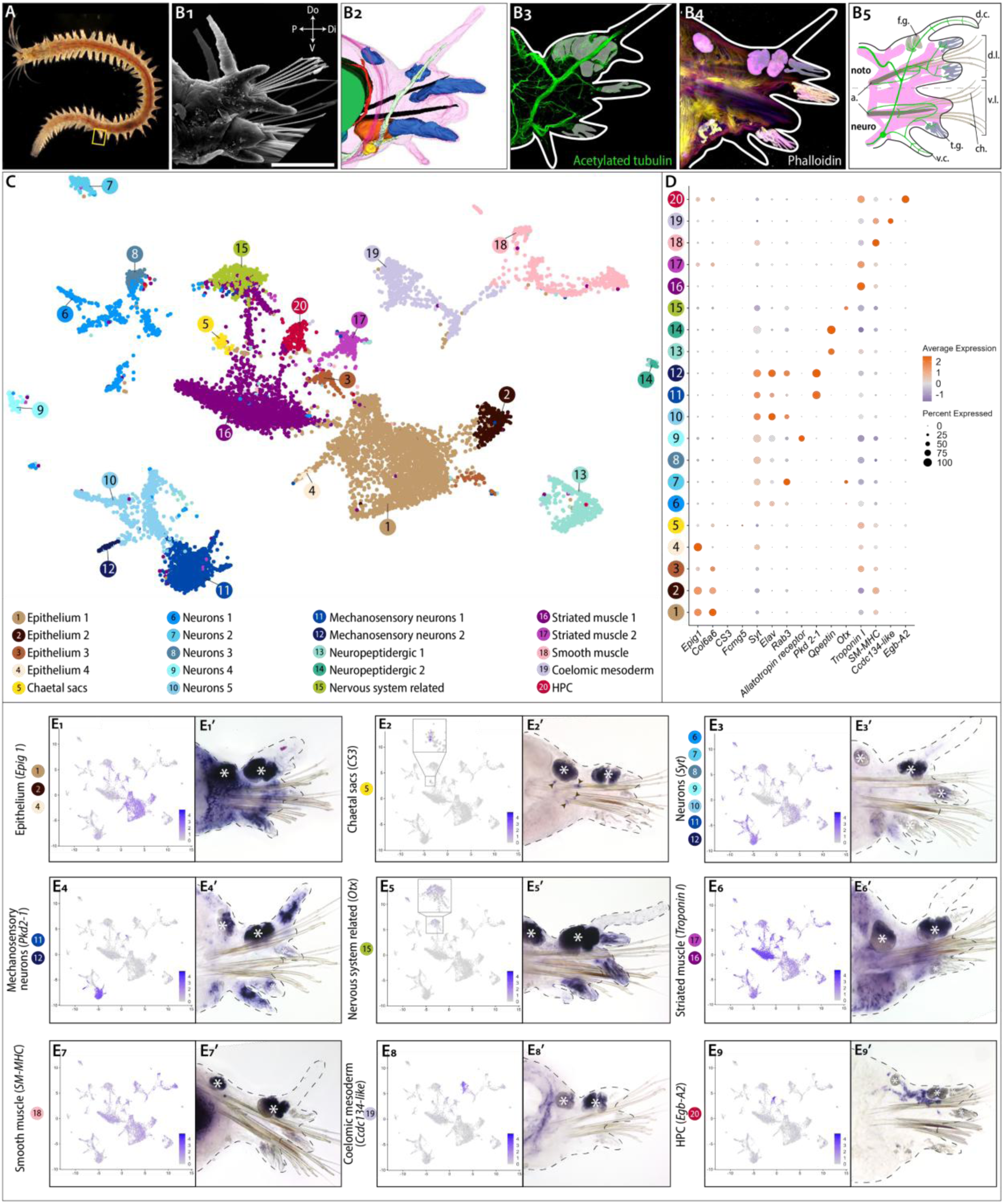
Tissue organization and cell diversity of *Platynereis’* parapodia. (A) Photo of a Platynereis juvenile worm. Each segment bears a pair of parapodia (yellow box). (B1-B4) Morphology and tissue organization of a parapodium, lateral views. (B1) Scanning electron microscopy (SEM) micrograph of a parapodium. (B2) 3D reconstruction of a parapodium made from stacks of semi-thin sections. Each parapodium bears a pair of aciculae (black) providing support to the structure. Glands (dark blue) and metanephridia (orange) are also present. Nervous projections from the ventral nerve chord forming a prominent parapodial ganglia (yellow) and innervating the cirri are also visible (pale green). (B3) Immunostaining against acetylated tubulin (green) labels the axon scaffold of the peripheral nerves present in a parapodium. (B4) Depth coded confocal z-stack projection of a phalloidin stained parapodium revealing the musculature. B3 and B4: A white line outlines the parapodium. Pale white inserts indicate the non-specific staining of the parapodial glands. (B5) Parapodium schematic representation with glands (grey and blue) and major nerves (green) and muscles (pink). noto. notopodium; neuro. neuropodium; a. acicula; f.g. flask-shaped gland, t.g. tubular-cell gland; d.l., dorsal lobe; d.c. dorsal cirri; v.l., ventral lobe; v.c. ventral cirri; ch. chaeta. (C) Uniform manifold approximation and projection (UMAP) of the annotated atlas with the 20 identified clusters colored according to their broad cell-population identity. (D) Dotplot of gene markers used to annotate the major cell populations in our dataset. (E1-E9) UMAP expression plot of marker genes for the major cell types. Cells expressing the gene of interest are colored purple. Cell populations overexpressing the gene in question are indicated on colored circles. (E1’-E9’) Whole Mount *In Situ* Hybridizations (WMISH) for the chosen gene markers for all clusters. Dark brown arrowheads, cells expressing *CS3*; Asterisks, glands. Do = Dorsal, V = Ventral, P = Proximal, Di = distal.

To obtain an in depth understanding of this complex structure, we assessed the cellular composition of *Platynereis* parapodia through a single cell RNA-seq approach. We used the ACME method (García-Castro et al., 2021) to obtain dissociated cells of individual parapodia, followed by 10X Genomics library preparation and paired-end Illumina sequencing. After assessing multiple quality control metrics, we retained 10 774 cells for downstream analysis, with an average of 5 023,5 reads/cell (Supp. Fig. 2A to C) which allowed us to obtain an overview of the cell composition for this structure which contains an average of 1 200 cells. Reads were then mapped against version 1.0 of the *Platynereis dumerilii* genome (Mutemi et al., 2025) (<u>GCA_026936325.1</u>). This procedure allowed us to robustly identify 20 transcriptionally distinct cell clusters of ectodermal and mesodermal nature that we considered as broad cell populations (Fig. 1C; Supp. Fig. 2D). We annotated all cell clusters, comprising epithelial, chaetal sacs, muscle, mesodermal, hematopoietic and neural cell populations by identifying known cell/tissue specific marker genes for *Platynereis* among their respective upregulated genes (Supp. Table 1, 2). We further validated most of these clusters by exploring the expression patterns of several gene markers on parapodia (Fig. 1D). We also extended their molecular signature by providing new upregulated gene markers (Supp. Table 3).

Among ectodermal tissues, we identified four clusters of epithelial cells, which collectively represent 30.6 % of the cells of this atlas. They are characterized by the specific overexpression of the epithelial markers *Epig1* and/or *Col6a6* (Epithelium 1 to 4, Fig. 1C, D and E_1_) (Stockinger et al., 2024), which was further confirmed by the expression pattern of *Epig1* in cells at the surface of the parapodia (Fig. 1E_1_’). To predict cell differentiation states, we looked at the expression of members of the Germline Multipotency Program (or GMP, a molecular signature of progenitor and/or stem cells in many animals, including *Platynereis* (Gazave et al., 2013; Juliano et al., 2010; Planques et al., 2019)), as well as known progenitor markers (Gazave et al., 2013a; Planques et al., 2019; Stockinger et al., 2024) and cell cycle genes (Demilly et al., 2013; Gazave et al., 2013b). This revealed that Epithelium 2 expresses many members of the GMP and the ectodermal progenitor marker *Sp/Btd* (Grimmel et al., 2016; Stockinger et al., 2024) (Supp. Fig. 3A). However, this population did not appear to highly express all the members of this specific set of markers (*i.e.*, progenitor gene module) in comparison to other cell populations (Supp. Fig. 3B). To better appreciate the diversity within those large epithelial cell populations, we subclustered cells contributing to Epithelium clusters 1 to 4, and chaetal sac cells into a dataset of 3 435 cells and generated a novel UMAP containing 12 clusters of epithelial identity (Supp. Fig. 4A-D, Supp. Table 4). To predict cell differentiation states and better infer the progenitor populations, we used the CytoTRACE package which infers the developmental potential of cells based on the diversity of genes expressed per cell and assuming that less differentiated cells have a higher diversity of gene expression (Gulati et al., 2020). Among the newly identified epithelial subpopulations, 8 of them (Epithelium 1a to 1h) are from the Epithelium 1 cluster subdivision. The majority of them express new gene markers of unknown expression pattern such as Epithelium 1c specifically expressing *Duox A* (Vullien et al., 2025). Epithelium 3 is sub-divided into two cell populations, one being *AchR alpha9-10+* (a marker of larval ciliated cells (Jékely et al., 2008). Epithelium 2 and 4 cell populations remain unchanged. Epithelium 2 overexpresses GMP genes, has a high CytoTRACE score, and may represent epithelial progenitors (Supp. Fig. 4A, C, D). Epithelium 4 specifically overexpresses the transcription factor *Tailless* (Achim et al., 2018) (Supp. Fig. 4A, B).

In addition to epithelial cells, ectodermal tissues contain discrete chaetal sacs cell populations (0.8 %) (Supp. Fig.2). Chaetal sacs are complex structures, at the origin of the worm chitineous bristles, composed of several cell types: a chaetoblast and its associated follicular cells (Gazave et al., 2017; Zakrzewski, 2011). While we were not able to discriminate between those two cell populations, as they are extremely scarce, we found a chaetal sac cluster as a whole, whose identity was determined thanks to the specific markers *Chitin Synthase 3 (CS3)* known to be expressed in the chaetoblasts (Gazave et al., 2017) and *Fcmg5* a marker of distal follicle cells (Zakrzewski, 2011) (Fig. 1C, D, E_2_,_2’_).

**Figure 2.**
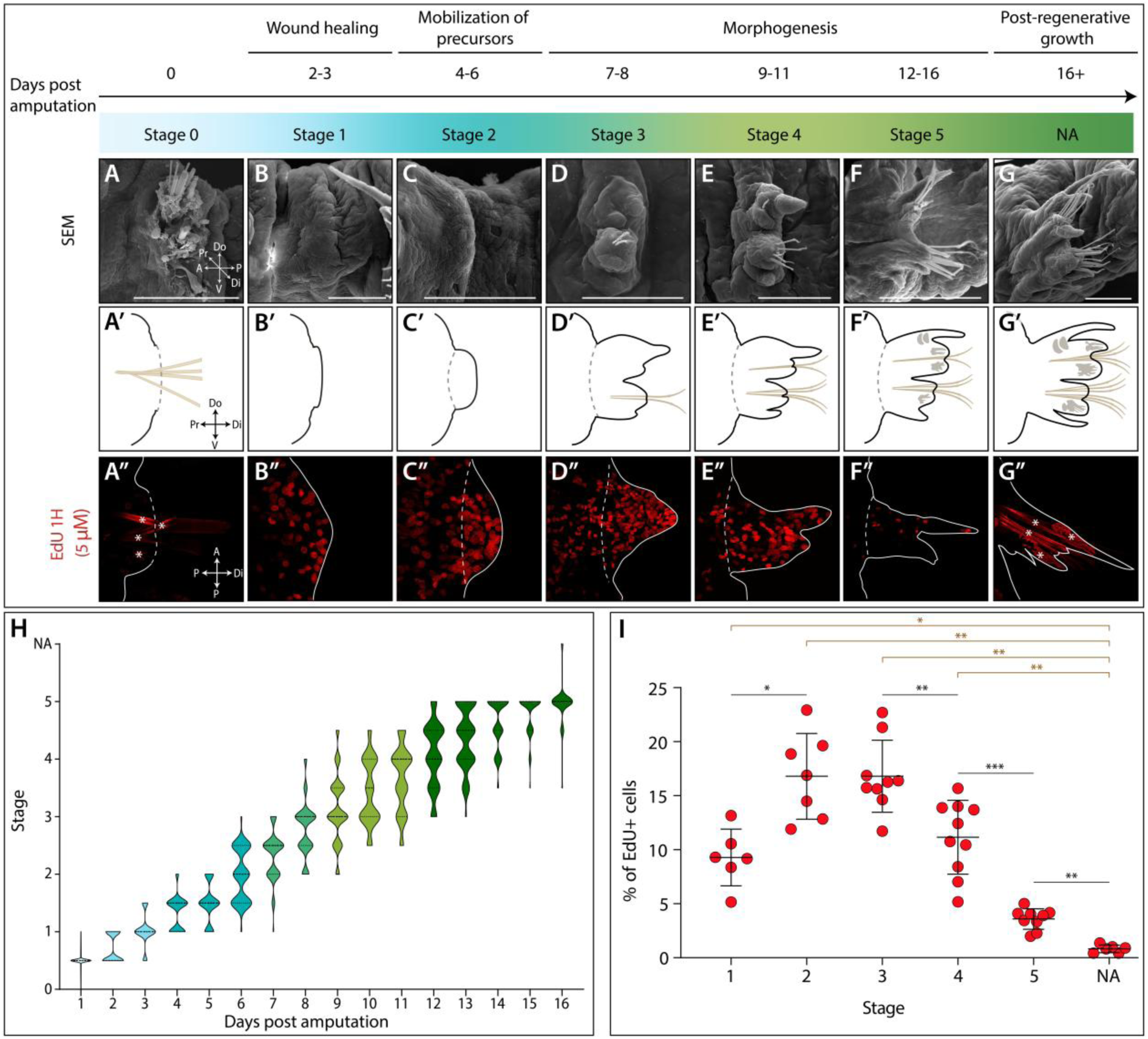
Morphological and cellular characterization of parapodial regeneration. (A-G) SEM micrographs of parapodial regeneration stages following a timeline and encompassing the three key steps of regeneration. Scale bar = 100 µm. (A’-G’) Schematic representation of the different regeneration stages. (A’’-G’’) EdU labelling (red) of the different regeneration stages after one hour of EdU incorporation (5 µM). White line, regenerate outline; white dotted line, amputation site; white asterisks, non-specific labelling of glands/chaetae. NA: non-amputated. (H) Color-coded violin plot depicting the distribution of the different regeneration stages reached by the worms (n=64) during 16 days scoring experiment. Dashed line, median; dotes lines, quartiles. (I) Proportion of EdU+ cells in the regenerating parapodia (n≥6) along the course of parapodial regeneration. Mean ± s.d. is indicated. *P< .05 ; **P< .01 ; ***P < .001 (Mann-Whitney test with Bonferroni-Dunn correction). Black, comparison between consecutive stages; brown, comparison to NA.

We also identified 10 nervous-system related clusters which collectively represent the majority of the cell types in the parapodia atlas (34.6 %) (Supp. Fig.2), as expected for a highly sensory structure (Fig. 1B_3_, C). Seven are neuron cells overexpressing the broad markers of differentiating (*Elav*) and differentiated neurons *Synaptotagmin* (or *Syt,* (Bideau et al., 2025; Denes et al., 2007)) (Fig. 1D, E_3_,_3’_). Among these neurons, two populations correspond to mechanosensory neurons, expressing the specific marker *Pkd2-1* (Fig. 1D, E_4_,_4’_, (Bezares-Calderón et al., 2018)). Additionally, we found two populations corresponding to neuropeptidergic cells (expressing *Qpeptin*, Fig. 1D, a *Platynereis* specific neuropeptide (Conzelmann et al., 2013)). The last one is considered as nervous system related thanks to its overexpression of *Otx,* a homeobox transcription factor previously described as involved in the patterning of embryonic neuronal tissues (Vopalensky et al., 2019) (Fig. 1D, E_5_,_5’_). All the neuronal populations identified show high scores for the previously described progenitor module (Supp. Fig. 3B), supporting their differentiated state, even though some overexpress specific members of the GMP signature, such as *Argonaute, Bruno A* and *B* (Supp. Fig. 3A, B). Neurons 2 and Neurons 5, along with Mechanosensory neurons 2 overexpress neuronal marker *Rab3* (Achim et al., 2018)) (Fig. 1D). Two other populations of neurons (1 and 3) are of unknown identity.

As for epithelial cells, to obtain a higher resolution of the parapodial nervous system, we reanalyzed only its cell populations (3 691 cells), generating a new clustering which allowed to identify a large diversity of neural cell types (n=21, Supp. Fig. 4E, Supp. Table 5). The Neurons 1 cluster appears to encompass a diversity of cell populations (Neurons 1a to e), all of them having a high CytoTRACE score but not overexpressing many GMP genes (Supp. Fig. 4E, G, H). Neurons 3 only contains two cell populations (Neurons 3a and b), with population 3b overexpressing *Achatin receptor 1* (Bauknecht & Jékely, 2015). The Neurons 2 cluster remains unchanged (Supp. Fig. 4G, H). Both have high CytoTRACE scores. The precise identity remains to be established for the majority of those neural cell populations, except for Neurons 1e which overexpress a Lophotrochozoa-specific neuropeptide *NKY-2*+ (Conzelmann et al., 2013). Neurons 1d specifically overexpress *Zic*, known to be expressed in the larval mushroom body neurons (Tomer, 2009). Neurons 4 is composed of two cell populations (a and b), 4a being *Gata1/2/3+* (a marker of multiple neuronal types including those in juvenile parapodial ganglia (Gillis et al., 2007)) and *Allatotropin receptor* 1+ (a neuropeptide receptor (Bauknecht & Jékely, 2015)). The central position of Neurons 1a, along with its projections toward populations 1b, 3a and 4b, and combined with its high CytoTRACE score suggests it represents a less differentiated/neural progenitor population. The large Neuron 5 population is sub-divided into four cell populations, with low CytoTRACE scores, including mechanosensory *NOMPC+* (Revilla-i-Domingo et al., 2021), photosensory *CNGAα+* and *Go-opsin+* (Ayers et al., 2018; Gühmann et al., 2015; Tosches et al., 2014), neuropeptidergic *EP+* and *FLamide+* (Conzelmann et al., 2013) neurons. Regarding the mechanosensory neurons (*Pkd2-1+*), the cluster number one comprises two cell populations (1a and 1b, *allatotropin+*) that are respectively *Post1+* and *Post1-*(a *Hox* gene family member (Kulakova et al., 2007)), while the number 2 is *Pkd1-1+* (another *Pkd* gene family member (Bezares-Calderón et al., 2018), Supp. Fig. 4E, F). Regarding the neuropeptidergic neurons, the population number one is divided into two cell populations (a and b), that express *Platynereis*-specific neuropeptide *Sll1+* (Conzelmann et al., 2013) and neuronal specification gene *Uncx4+* (Achim et al., 2018), respectively, while the Neuropeptidergic 2 cells overexpress *Platynereis*-specific neuropeptide *SHM+* (Conzelmann et al., 2013). The nervous system related cluster (*Otx+*) remains unchanged.

Finally, we investigated the mesodermal tissues and identified three populations of muscles, representing 26.4 % of the atlas (Fig. 1C, D; Supp. Fig. 2). Two of them are striated muscles, both overexpressing the striated muscle marker *Troponin I*, showing low progenitor module scores and a moderate overexpression of GMP genes, supporting the idea that those cells are fully differentiated (Fig. 1D, E_6_, E_6’_; Supp. Fig. 3A, B). The third one corresponds to smooth muscles, overexpresses *SM-MHC* (smooth-muscle myosin heavy chain, (Brunet et al., 2016), Fig. 1D), has a low progenitor module score despite highly expressing some GMP genes (Fig. 1D, E_7_, E_7’_; Supp. Fig. 3A, B). 5.7 % of the atlas corresponds to cells of the coelomic mesoderm, specifically overexpressing *Ccdc134-like* (Stockinger et al., 2024) (Fig. 1E_8_, E_8_’). Those cells may be progenitor cells, as they have high progenitor module scores, highly overexpress GMP, as well as cell cycle genes (*PCNA, cyclins*, and others (Gazave et al., 2013)), and mesodermal progenitor markers (*P2x, Chd3/4/5b, Prrx* (Planques et al., 2021; Stockinger et al., 2024)) (Supp. Fig. 3A, B). Our clustering also allowed us to identify the discrete hemoglobin-producing cells population (HPC, 1.8%, Supp. Fig. 2) thanks to the specific expression of several extracellular globins (*Egb-A2, A1a, A1b* (Song et al., 2020)) (Fig. 1C, D, E_9_, E_9’_).

To assess more precisely the diversity of mesodermal cell populations, we extracted them (n= 3 648 cells) and performed a subclustering that allows the identification of 11 cell populations (Supp. Fig. 4I, J; Supp. Table 6). The Striated muscle 2 and HPC clusters remain unchanged. The Striated muscle 1 is subdivided into three distinct cell populations, 1a and 1c are *Mef2+* (a myogenic transcription factor (Brunet et al., 2016)). Furthermore, based on the overexpression of several GMP genes, population 1c might represent striated muscle progenitors despite its low CytoTRACE score (Supp. Fig. 4K, L). The Smooth muscle cluster comprises four distinct sub populations of unknown identity with high CytoTRACE scores. Smooth muscle b also overexpresses many GMP and cell cycle genes and may represent smooth muscles progenitors (Supp. Fig. 4K, L). The coelomic mesoderm comprises two cell populations (a and b), with population b overexpressing GMP and cell cycle genes, and with a high CytoTRACE score suggesting it might represent smooth muscle progenitors (Supp. Fig. 4K, L).

Overall, our single-cell atlas has likely uncovered the majority of the cell population diversity present in *Platynereis* parapodia (with the notable exception of metanephridia, which we were unable to identify). Consequently, we identified both differentiated cells and putative progenitor cells, the latter of which may be implicated in tissue homeostasis and/ or regeneration. The integration of morphological and tissue data into this comprehensive cell transcriptomic atlas allows for a unique and deep understanding of this major annelid complex structure.

### *Platynereis* parapodia regeneration: morphological, cellular and molecular insights

Following upon this new knowledge of *Platynereis’* parapodia, we then assessed how they regenerate. Hence, we performed a thorough study of this process at the morphological, cellular and molecular levels.

#### Morphological and cellular characterizations of parapodial regeneration

We performed amputations of a single parapodium located in the 40^th^ posterior segment along the trunk of 5-months juvenile worms. Using two complementary approaches, scanning electron microscopy (SEM), which allows for fine descriptions of the external morphology, and bright field microscopy, we defined 6 regeneration stages that follow the typical steps of regeneration (Fig. 2A to G, H). Immediately upon amputation, at stage 0 (0 day post amputation or dpa), the tissues are damaged, pieces of bristles are still attached to the structure (Fig. 2A, A’). Between 2 to 3 dpa, at stage 1, a wound epithelium is reformed providing a physical barrier between the damaged structure and its environment, this is the wound healing step of regeneration (Fig. 2B, B’). Between 4 to 6 dpa, at stage 2, a mass of cells under the wound epithelium appears. This corresponds to the mobilization of precursor cells participating in this regeneration step (Fig. 2C, C’). From 7 to 16 dpa, there is a long phase of morphogenesis until the complete reformation of the amputated structure (Fig. 2D to G’). At stage 3 (7 to 8 dpa), the notopodium with a small dorsal cirrus and the neuropodium, bearing few chaetae, start to be distinguishable (Fig. 2D, D’). At stage 4 (9 to 11 dpa), while still not fully reformed, both lobes bear chaetae and their respective cirri (Fig. 2E, E’). At stage 5 (from 12 to 16 dpa), the dorsal glands are reformed, notopodium and neuropodium increase in size (Fig. 2F, F’). This latest stage is followed by a phase of post-regenerative growth until the parapodium becomes indistinguishable from neighboring ones (Fig. 2G, G’). Scoring 64 worms daily for 16 days confirmed that these stages are reproducible, with most worms reaching them at a similar pace. However, a certain degree of variability among the worms was also observed (Fig. 2H, Supp. Table 7).

In many regenerative species, including *Platynereis*, cell proliferation is a mandatory actor of regeneration success (Planques et al., 2019). We thus determined the cell proliferation dynamics during the course of parapodial regeneration through EdU labelling (which marks all cells passing through S phase) and expression of the cell cycle gene *PCNA* (Fig. 2A’’ to G’’, I; Supp. Fig. S5A to A_6_; Supp. Table 8). Although cell proliferation is quasi-inexistent in the tissue abutting the amputation plane at stage 0, at stage 1, many EdU+ cells are found (about 10 %), both close to the amputation site and in the non-amputated tissues of the trunk. This proportion of EdU+ cells significantly increases at stages 2 and 3, reaching about 17 % of the cell mass formed. After stage 3, the proportion of EdU+ cells drastically reduces and in fully regenerated/non-amputated parapodia, the proportion of proliferative cells is negligible (less than 1 %). This dynamic is confirmed by the expression pattern of *PCNA* whose expression is observed as early as stage 1, is large at stage 2 and progressively diminishes during morphogenesis (Supp. Fig. 5A to A_6_). Although cell proliferation appears to be important for parapodial regeneration, we tested whether it is mandatory by using the anti-proliferative agent hydroxyurea (HU) and analyzing its effect on regeneration success (Supp. Fig. 5B to F’). Thus, we performed HU treatment starting at different time points after amputation (from 0 dpa to 9 dpa) and pursued treatment until 16 dpa, scoring the stages reached by worms at 2, 5, 7, 9, 12 and 16 dpa (Supp. Fig. 5B, C, Supp. Table 9). We observed that in the presence of HU since 0 dpa, regeneration is stopped at stage 1.5 and only a small cell mass is formed. We also noted that all worms whose HU treatment started at 0 and 2 dpa did not reach stage 2. We confirmed that HU did indeed block cell proliferation, by performing an EdU pulse at 5 dpa followed by a chase phase in HU (Supp. Fig. 5D). As expected, in the presence of HU, cells are blocked in S phase. This is illustrated by their large and homogenous EdU nuclei labelling, in comparison to the stippled one – obtained through cell divisions – for the control worms (Supp. Fig. 5E to F’). Hence, this set of experiments show that cell proliferation is mandatory for regeneration to proceed past stage 2.

Altogether, we showed that parapodial regeneration is a reproducible process, following 6 stages encompassing the key regeneration steps. This process relies on the formation of a proliferative mass of cells that we call blastema. Cell proliferation appears mandatory upon blastema formation, for blastema growth and for parapodial morphogenesis.

#### Kinetics of structure reformation during regeneration

To further characterize *Platynereis* parapodial regeneration, we studied the kinetics of reformation of the different parapodial structures and tissues. To this end, we investigated the expression patterns of 13 genes previously shown to be involved in appendage, muscle, hematopoietic and nervous system patterning and differentiation, and fluorescently labelled differentiated muscles and axon projections (Fig. 3; Supp. Fig. 6) (Planques et al., 2019). Those results are summarized below and schematically represented in Fig. 3.

**Figure 3.**
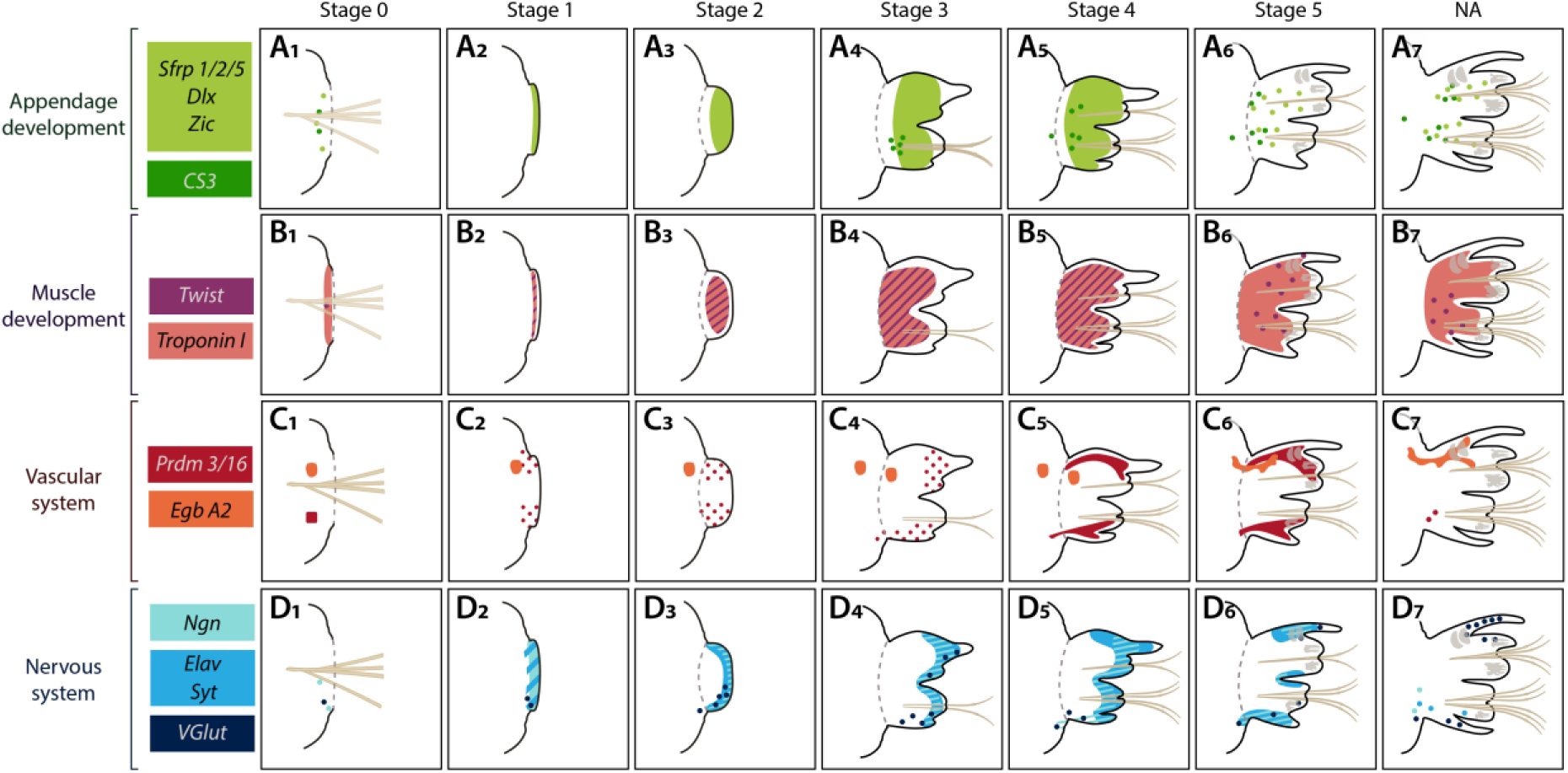
Summary of the expression patterns of genes involved in organ or tissue patterning, and differentiation, during parapodia regeneration. (A to D) Schematic drawings of the regenerating parapodia at stages 0 to 5 and of non-amputated (NA) parapodia. The territories of expression of various organ, structure, and tissue marker genes are shown and color-coded. Grey dotted line: amputation plane.

We first investigated appendage formation with *Dlx, Zic* and *Sfrp 1/2/5* known to be expressed in the parapodia primordia (Grimmelikhuijzen et al., 1989; Planques et al., 2019), and *CS3*, the chaetoblast marker indicating an advanced differentiation of the structure (Fig. 3A_1_ to A_7_; Supp. Fig. S6A_1_ to D_7_). *Dlx, Zic* and *Sfrp1/2/5* are expressed as early as stage 1 in the wound epithelium and then more largely in the blastema at stages 2 and 3. Their expression is progressively restricted, and at stage 5, is similar to the non-amputated (NA) control. In contrast, *CS3* starts to be expressed later, at stage 3 when chaetae are reformed and at stage 5 its expression is as for NA.

We also studied the formation during regeneration of mesodermal derivatives, namely muscles (Fig. 3B_1_ to B_7_; Supp. Fig. S6E_1_ to G_7_), blood vessels, and hemoglobin producing cells (HPC) (Fig. 3C_1_ to C_7_; Supp. Fig. S6H_1_ to I_7_). *Twist*, a marker of somatic muscle development (Pfeifer et al., 2013), is broadly expressed in mesodermal tissues from stage 1 until stage 4, then its expression fades away. *Troponin I*, a specific marker of striated muscles (Brunet et al., 2016) is first expressed in the blastema (stage 2), and its expression is maintained until the end of the process. This is confirmed by the Phalloidin labelling of actin fibers which reveals a retraction of the muscle fibers during the wound healing step (stage 1), while they progressively infiltrate the blastema and regenerated structure later on (Supp. Fig. 6G_1_-G_7_). Blood vessel marker *Prdm3/16* (Planques et al., 2019) and extracellular globin *Egb A2* (Song et al., 2020) are both expressed early upon amputation (stage 1), in discrete cells that will progressively form the vessels and its associated HPC cells in the lobes.

We then studied, throughout regeneration, the expression patterns of a set of markers known to be involved in the formation of *Platynereis* nervous system and hence constituting a neurogenic cascade (Bideau et al., 2025; Denes et al., 2007; Kerner et al., 2009) (Fig. 3D_1_ to D_7_; Supp. Fig. S6K_1_ to N_7_). The neural progenitor marker *Neurogenin (Ngn)* is largely expressed at stage 1, while its expression is moderate in the blastema at stages 2 and 3. During morphogenesis, it is mainly found in cells in the cirri. *Elav,* a marker of differentiating neurons, is also expressed since stage 1, while its expression increases during blastema formation and early morphogenesis. At late morphogenesis stages, its expression fades away. A similar expression pattern dynamic is observed for *Synaptotagmin (Syt*), a broad marker of differentiated neurons. The glutamatergic neuron marker *VGluT* is expressed in few cells of the stage 2 blastema, and its expression is restricted to scattered cells in the lobes and cirri during morphogenesis. Neurite labelling with acetylated tubulin revealed that upon amputation and up to stage 1, the nerves are a bit disorganized (Supp. Fig. S6N_2_, white arrow) but very rapidly, they extend their projections to innervate the blastema (stage 2) and all the parapodial structures during morphogenesis, notably reforming the nervous projections found in the parapodial cirri (Supp. Fig. 6N_1_ to N_7_).

This expression pattern analysis has yielded a comprehensive understanding of the kinetics of the reformation of multiple parapodial structures and has indicated that the stage 2 blastema may contain both undifferentiated progenitor/ stem cells and differentiated cell types.

### Parapodial blastema atlas reveals progenitor populations and differentiation trajectories for the main tissue lineages

We thus investigated more specifically this key blastema stage and performed, as for non-amputated parapodia, scRNA-seq on dissociated cells of about 800 individual blastemas (Fig. 4). After assessing the same control metrics as before, we retained 3 801 cells for downstream analysis with an average of 6 459.1 reads/cell (Supp. Fig. 7A to C) which allowed us to study in depth this structure comprised of an average of 500 cells. A total of 19 transcriptionally distinct cell clusters were identified. Automatic annotation transfer using the NA parapodia dataset as a reference was done to assign broad tissue categories to each cluster, and refined using known gene markers as before (Fig. 4A, B, Supp. Table 10). We also computed dot plots for GMP signature genes, ectodermal and mesodermal progenitors, as well as cell cycle genes, and progenitor module scores (Fig. 4C, D). Finally, we performed WMISH for some of those genes (Supp. Fig. 5A-A_6_, 8).

**Figure 4.**
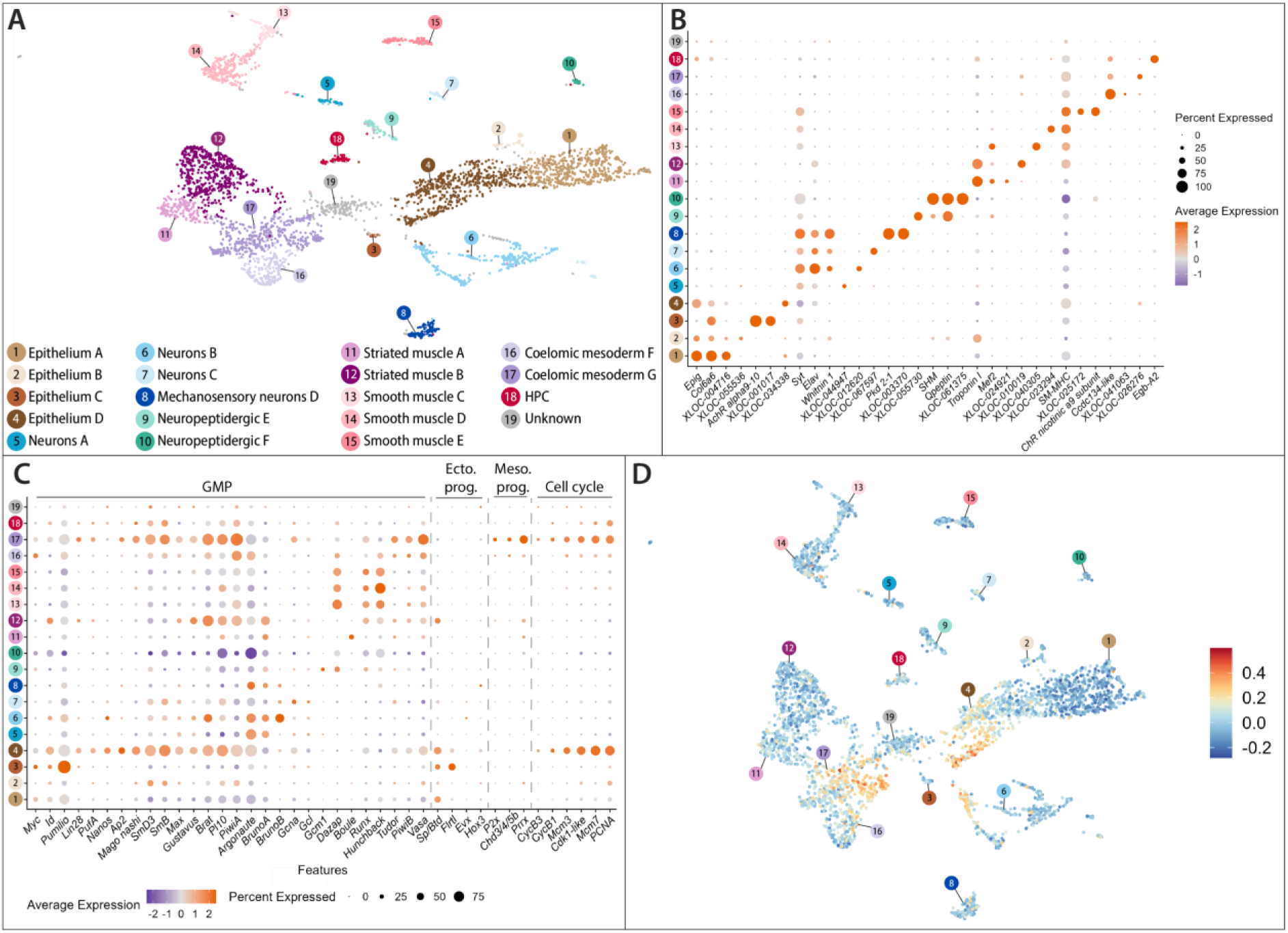
Cell diversity in the parapodia blastema. (A) UMAP of the annotated parapodia blastema atlas with the 19 identified clusters colored according to their cell type identity. (B) Dotplot of known and new gene markers for the identified cell populations. (C) Dotplot of genes belonging to the GMP signature, markers of ectodermal and mesodermal progenitors, and cell cycle. (D) UMAP visualization of the progenitor module expression score per cell. High scores (red) indicate cells with high expression of the gene module of interest.

We identified four populations of epithelial cells, which collectively correspond to 29.4 % of the blastema atlas. Three of them (Epithelium A, B and D) specifically overexpress *Epig1* and *Col6a6* and, based on the annotation transfer, appear to share their molecular identities with epithelial populations previously identified in the non-amputated parapodia (Fig. 4A, B). A large Epithelium A cluster (13.9 %) corresponds transcriptionally to the populations Epithelium 1a to g found in the parapodia atlas and is enriched for the *Sp/Btd* ectodermal progenitor marker. Epithelium B does not overexpress any ectodermal progenitor markers and is transcriptomically similar to Epithelium 1a and 1d. Epithelium A and B are mostly not enriched for GMP nor cell cycle genes. In contrast, Epithelium D is enriched for GMP signature genes (Supp. Fig. 8A_1_ to A_6_) and ectodermal progenitor markers (*Sp/Btd* and *Flrtl -* the latter of which we confirmed to be expressed in epithelial cells (Fig. 8A_7_)), appears highly proliferative (based on the overexpression of all tested cell cycle genes), and has a high expression of our predefined gene module (Fig. 4C, D). Epithelium D is large (13.5 %), and shares a transcriptomic identity with Epithelium 1a, 1c, 1d, 1e and 2 from the parapodia atlas. A fourth small cell population (Epithelium C) specifically overexpresses two ectodermal progenitor markers, does not overexpress GMP genes (with the exception of *Pumilio*), and does not appear proliferative based on its gene expression (Fig. 4D). Altogether this is suggestive of a mix of epithelial cell populations at different levels of differentiation, with Epithelium D as a potential broad epithelial progenitor population.

To extend our cell differentiation hypothesis, we performed trajectory analysis with Slingshot on the blastema dataset (Fig. 5, Supp. Fig. S9). To this end, epithelial cells were extracted and sub-clustered, allowing to discriminate 7 cell populations. Six of them correspond to the previously defined Epithelium A, B (now divided in B1 and B2), C and D (now separated in D1 and D2) populations (Fig. 4A, B) and one newly identified cell population (Epithelium E) sharing a molecular signature with Epithelium A and D (Fig. 5A, Supp. Table 11). The cell trajectories were charted on the UMAP representation. The starting point was chosen as Epithelium D2 based on its CytoTRACE score and GMP signature expression (Supp. Fig. 9A-B). Four differentiation trajectories were suggested and showed a lineage progression from D2 through potentially more engaged intermediate progenitors D1 and/or E which may give rise to more differentiated A, B1, B2 and C epithelial cell populations (Fig. 5A, Supp. Fig. 9C-F).

**Figure 5.**
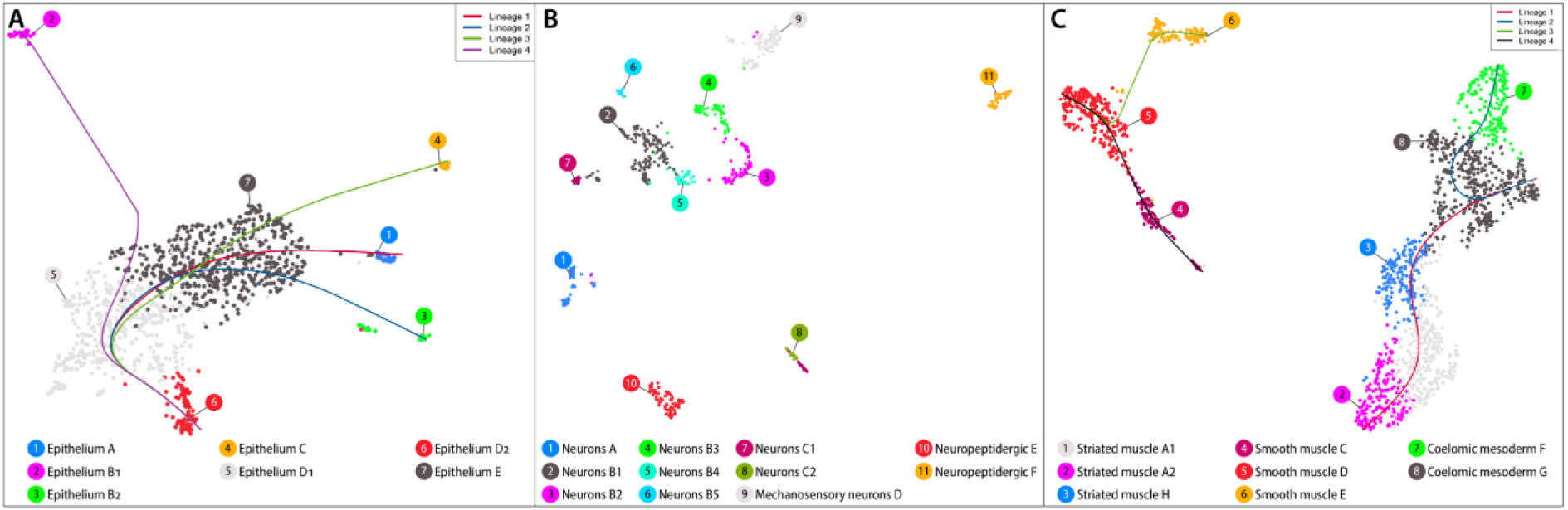
Parapodia blastema cell subpopulations per broad cell category and associated cell trajectory analysis. Subsets of epithelial (A), neural (B), and mesodermal (C) parapodia blastema cell populations. For the epithelial and mesodermal populations, differentiation lineages are indicated as lines on their respective UMAPs.

We also identified 6 nervous-system related clusters (16.4 % of the atlas, (Fig. 4A, B)). Three populations of neurons (A to C) overexpress *Syt, Elav*, and/or *Rab3* while the mechanosensory neurons D and the populations of Neuropeptidergic cells E and F are enriched in *Pkd2-1* and *Qpeptin*, respectively. None of them are enriched in GMP members although some cells have high expression of our progenitor gene module, supporting their differentiated state, except for Neurons B (Fig. 4C, D). We extracted and sub-clustered all the neuronal cells into 11 cell populations. A, D, E and F remain unchanged, Neurons C is divided into two cell populations (C1 and C2) and Neurons B is separated into 5 cell populations (B1 to B5) (Fig. 5B, Supp. Table 12). Among them, B2 overexpresses GMP markers and no cell cycle genes (Supp. Fig. 9G, H), but very few of its cells express known neural progenitor markers *Ngn*, and *NeuroD* (Simionato et al., 2008). We tried to infer cell trajectories among these nervous system populations using Neurons 2 as the starting point. However, the results were inconclusive (not shown), supporting the idea that the differentiation of the nervous system might be too advanced at this regeneration stage to correctly infer the trajectories at play.

Finally, we found 8 cell populations of mesodermal tissues which collectively represent 47.6 % of the blastema atlas: two populations of striated muscles (overexpressing *Troponin I*), three populations of smooth muscles (overexpressing *SM-MHC*), two populations of coelomic mesoderm (overexpressing *Ccdc134-like*), and an HPC population (overexpressing *Egb-A2*) (Fig. 4A, B). The latter may correspond to cells from non-amputated tissues (inevitably recovered during the collection procedure) that may contaminate the blastema sample as globin markers are known to be expressed in more differentiated tissues (Fig. 3, Supp. Fig. 6I_1_-I_7_; (Song et al., 2020)). Striated muscles A and B partly share a molecular signature with Striated muscles 1 from non-amputated parapodia. They both have a low progenitor module scores, but Striated muscles B overexpresses some GMP markers and may contain not fully differentiated cells (Fig. 4C, D). Smooth muscles C, D and E appear as relatively differentiated cells. Among the coelomic mesoderm populations, G overexpresses many GMP and cell cycles genes, as well as mesodermal progenitors (such as *Prrx*, Supp. Fig. 8A_8_), has a high progenitor module score, and hence probably corresponds to the coelomic mesoderm progenitor population (Fig. 4C, D). We sub-clustered all muscles and coelomic mesoderm cells from the blastema dataset and mainly identified the same populations, except for the striated muscles (Fig. 5C, Supp. Table 13). The Coelomic mesoderm G was fixed as one starting point for some trajectories based on its CytoTRACE scores and expression of GMP genes, progenitor markers, and cell cycle genes (Supp. Fig. 9 I, J). It appears to give rise to a less proliferative and GMP-enriched Coelomic mesoderm F, but also to the differentiated striated muscles populations A (A1 and A2) through an intermediate population of striated muscles (H) (Fig. 5C, Supp. Fig. 9K, L). The other starting point was fixed as Smooth muscle D, which appears to give rise to both Smooth muscle C and E (Supp. Fig. 9M, N). All smooth muscle populations are already well differentiated and disconnected from the other mesodermal populations (Fig. 5B, Supp. Fig. 9I, J).

Thus, thanks to this single cell data set, we unraveled the precise cell type composition of the parapodia blastema and identified potential progenitor populations for the main tissues which allowed us to determine partial cell trajectories.

### Blastema comparison highlights common features for parapodial and posterior successful regeneration in *Platynereis*

In addition to parapodial regeneration, *Platynereis* worms have the ability to successfully regenerate their complex posterior part, a process relatively well known at the morphological, cellular and molecular levels (Bideau et al., 2024, 2025; Kostyuchenko et al., 2019; Paré et al., 2023; Planques et al., 2019; Stockinger et al., 2024; Vervoort & Gazave, 2022; Vullien et al., 2025). Aiming to assess successful regeneration events in *Platynereis*, we compared parapodial and posterior regenerations at the single cell level, focusing more specifically on the similar key blastema stage (*i.e.* mass of cells with no visible morphogenesis showing a pic of cell proliferation and large expressions of the GMP markers (Planques et al., 2019)).

#### A detailed posterior regeneration blastema atlas

We first generated a posterior blastema dataset using the same methodology as for the parapodial blastema sample. We then integrated this dataset with public single-cell data, at the same stage, from the literature (Stockinger et al., 2024) to produce a very large dataset, which we analyzed in depth (Fig. 6).

**Figure 6.**
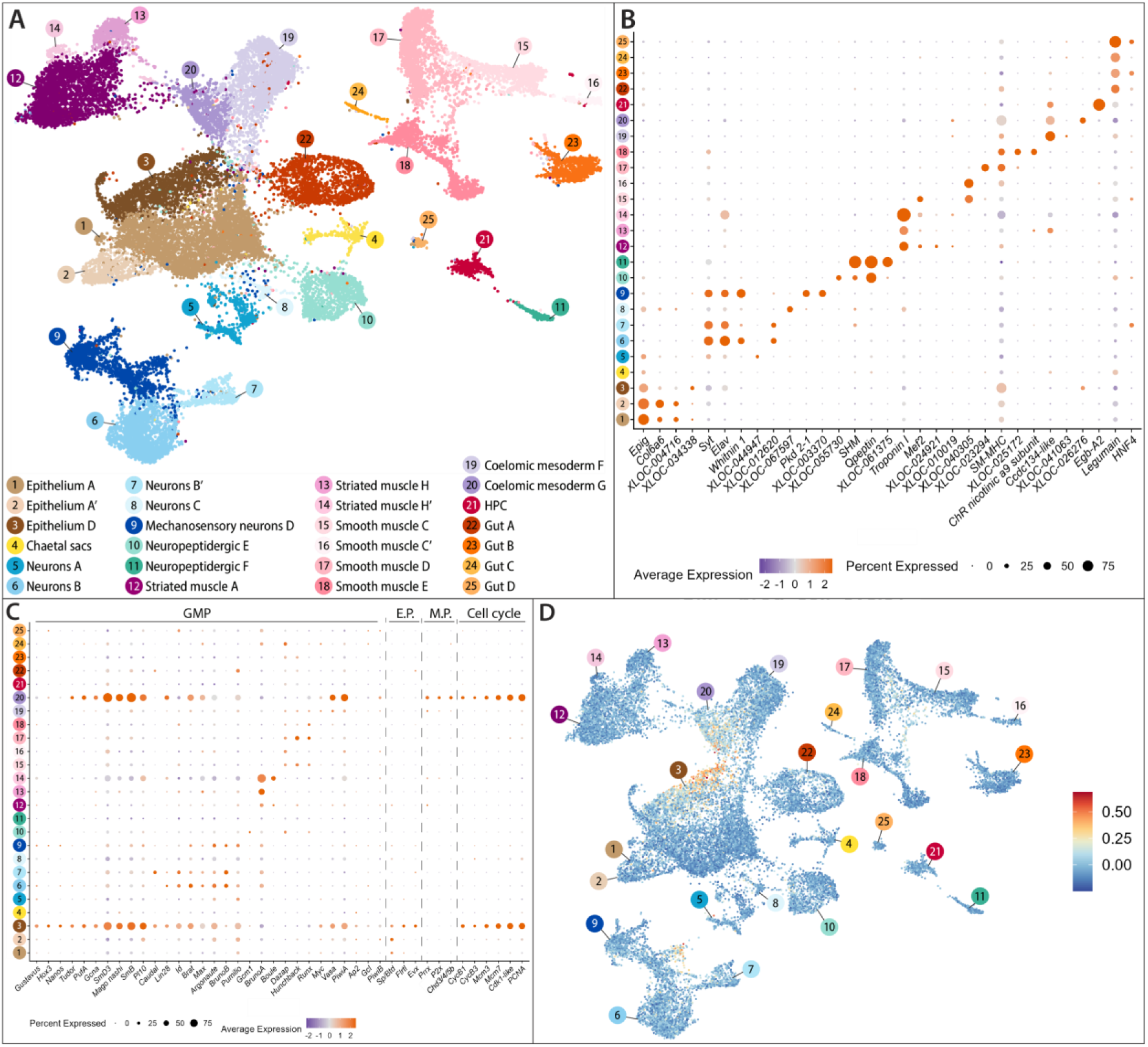
Cell diversity in the posterior part blastema. (A) UMAP of the annotated posterior part blastema atlas with the 25 identified clusters colored according to their cell population identity. (B) Dotplot of known and new gene markers for the identified cell populations. (C) Dotplot of genes belonging to the GMP signature, markers of ectodermal and mesodermal progenitors, and cell cycle. (D) UMAP visualization of the progenitor module expression score per cell. High scores (red) indicate cells with high expression of the gene module of interest.

After the quality check, we obtained 28 053 cells with an average 3 121.5 reads per cell. A total of 25 transcriptionally distinct cell clusters were identified (Fig. 6A). Automatic annotation transfer using the parapodia blastema dataset as a reference was done to assign broad tissue categories to each cluster which was then narrowed down by looking at specific markers for each population (Supp. Table. 14). Annotations were further clarified by manually assessing the overexpression of key markers (Fig. 6B, Supp. Table 14, 15). When possible, a similar nomenclature for the cell populations was used between parapodial and posterior blastema atlases for an easier comparison. As for the parapodial blastema atlas, we also computed progenitor module scores, provided dot plots for GMP signature genes, ectodermal and mesodermal progenitors, as well as cell cycling genes (Fig. 6C, D; Supp. Fig. 10A-D). We identified three populations of epithelial cells (A, A’ and D), which collectively correspond to 24 % of the posterior blastema atlas (Fig. 6A; Supp. Fig. 10D). Epithelium D, which overexpresses GMP, cell cycle genes and ectodermal progenitor markers, may correspond to a broad epithelial progenitor population, reflected in its high module score (Fig. 6C, D). A small population of chaetal sacs cells is also found (1.4%) and may correspond to cells from non-amputated tissues. After extraction and subclustering of epithelial cells, eight populations were determined, four of them were identified as sub-populations of A, two of D, and two new populations were found (Fig. 7A, Supp. Table 16). Epithelium D2 (with a high CytoTRACE score, overexpressing GMP and cell cycle genes, Supp. Fig. 11A_1_, A_2_) was chosen as the trajectories’ starting point and two trajectories were proposed (Fig. 7A). Trajectory one links D2 to A (A1 to A3) through D and F (Supp. Fig. 11A_3_). Trajectory two ends in Epithelium G through D (Supp. Fig. 11A_4_).

**Figure 7.**
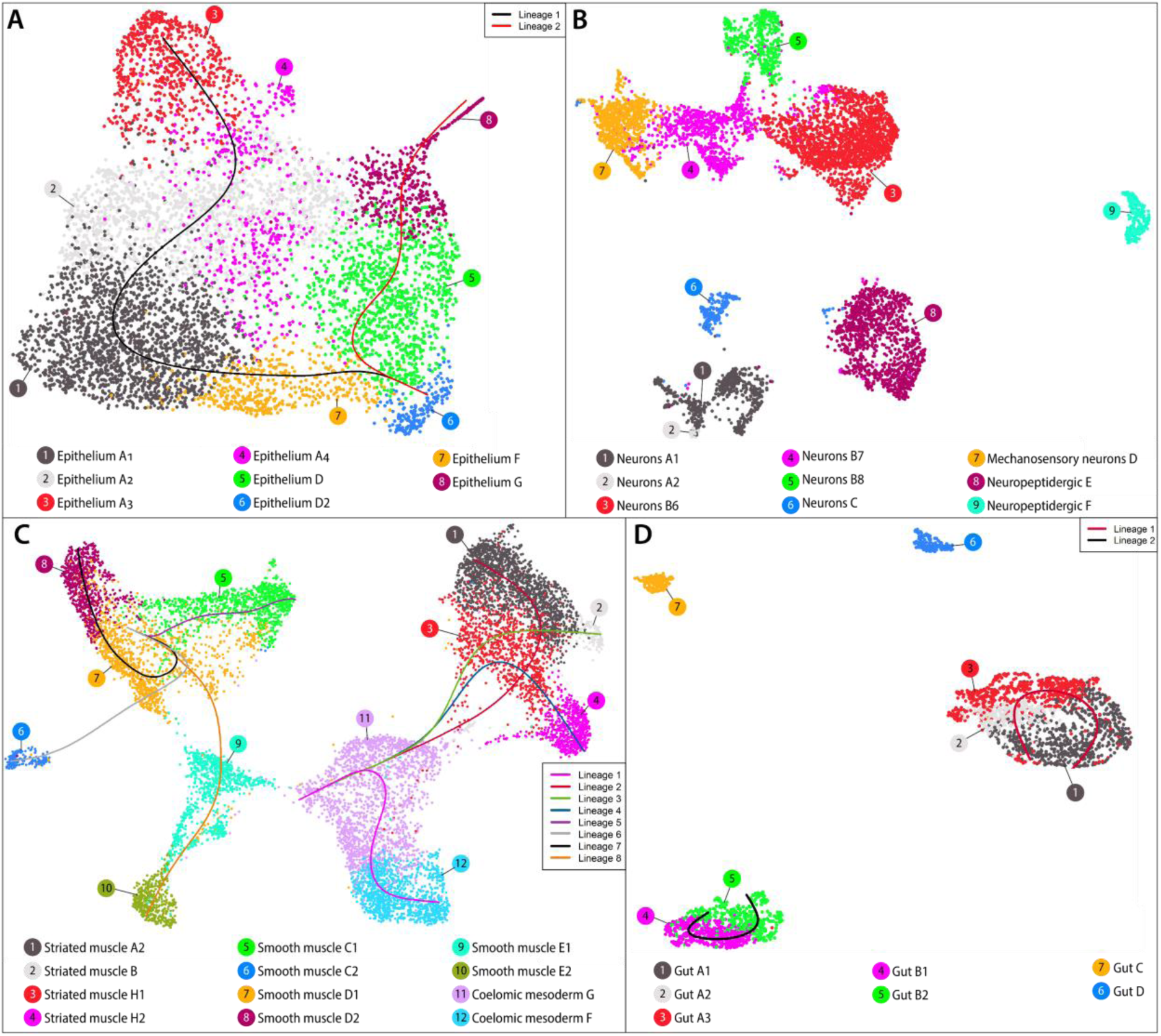
Posterior blastema cell subpopulations per broad cell category and associated cell trajectory analysis. Subsets of epithelial (A), neural (B), mesodermal (C), and gut (D) posterior blastema cell populations. For the epithelial, mesodermal, and gut populations, differentiation lineages are indicated as lines on their respective UMAPs.

Seven populations of neural cells were also identified (22.8 % of all cells, Fig. 6A, B; Supp. Fig. 10D). Four of them were classified as neurons (A, B and B’, C, and Mechanosensory neurons D), with Neurons B and B’ overexpressing GMP genes in a few cells (Fig. 6C, D). Two populations of neuropeptidergic differentiated cells (E and F) were also found. After extraction and subclustering of neural cells, we discriminated 9 populations (Fig. 7B, Supp. Table 17). The Neurons C, Mechanosensory neurons D and neuropeptidergic cells populations remained the same. Neurons A and B are divided into several subpopulations (2 and 3 respectively). The central population of Neurons B7 (with high CytoTRACE score,overexpressing GMP genes, and with a few cells overexpressing known neural progenitor markers, Supp. Fig. 11B_1_, B_2_) was chosen as the trajectories’ starting point. However, as for the parapodia blastema, we failed to determine conclusive cell trajectories (results not shown).

This posterior blastema atlas also contains populations of mesodermal cells (40%, Fig. 6A, B; Supp. Fig. 10D) including three populations of striated muscles (A, H, and H’), four populations of smooth muscles (C, C’, D and E) and two populations of coelomic mesoderm (F and G). Furthermore, as the population of coelomic mesoderm G overexpresses GMP, cell cycle genes and mesodermal progenitor markers, and has a high module score, it may correspond to a broad mesodermal progenitor population (Fig. 6C, D). A small population of HPC potentially coming from non-amputated tissue cells is also found (1.5%). Specific clustering of mesodermal cells allowed us to identify 12 cell populations (Fig. 7C, Supp. Table 18). Using the same GMP and CytoTRACE score criteria as before, Coelomic mesoderm G and Smooth muscle D1 were selected as starting points and eight trajectories were proposed (Supp. Fig. 11C_1,2_). The Coelomic mesoderm G population may give rise to Coelomic mesoderm F and to Striated muscles A2, B and H2 through H1 (as mixed population of A and B) (Supp. Fig. 11C_3-5_). Four other trajectories link Smooth muscles D1 directly to C1, C2 and D2, and to E2 through E1 (Supp. Fig. 11C_6-10_).

Finally, we found 4 populations of gut cells (A to D, for a total of around 10%), all of them overexpressing the *legumain protease precursor*, a digestive enzyme (Williams et al., 2015). Additionally, populations A, B, and D overexpress midgut-specific transcription factor *HNF4* (Achim et al., 2018). New clustering focused on gut cells allowed to discriminate 7 populations (Fig 7D, Supp. Table 19). Although C and D remained the same, A and B are divided into 3 and 2 sub-populations, respectively. Based on CytoTRACE scores and GMP marker expression (Supp. Fig. 11D_1,2_), populations A1 and B1 were selected as the starting populations, leading to two different trajectories. In the first one, Gut A1 gives rise to Gut A3 and later on to Gut A2 (Supp. Fig. 11D_3_). In the second, Gut B1 cells give rise to Gut B2 cells (Supp. Fig. 11D_4_). Both Gut C and D appear to be disconnected from the other gut populations (Fig. 7D, Supp. Fig. 11D_3,4_).

#### Comparative analysis of parapodial and posterior blastema cell diversity and cell trajectories

The detailed cell atlas obtained for both blastemas enables a fine-grained comparison of the cell populations distribution between the two structures (Fig. 8A). On a global scale, the cell population diversity and proportions are similar but noticeable exceptions were identified. Overall, the parapodia blastema contains more epithelial cells (29.4 % *versus* 24 % for the posterior part blastema) and also presents two specific and discrete cell populations absent from the posterior part (epitheliums B and C). The neural cell types are also very comparable between the two structures. Neurons A, B and C represent similar proportions of cells of the structure, while the mechanosensory neurons and neuropeptidergic cells appear to be in higher proportions in the posterior part blastema (2.58 % *versus* 5.98 %). Regarding the mesodermal cell populations, the parapodial blastema shows a higher proportion of coelomic mesoderm populations (15.75 % *versus* 10.91 %) and contains more G – that may represent broad progenitors - than F, while the opposite is true for the posterior part. The populations and proportions of smooth and striated muscles are more or less the same with the exception of an additional striated muscle population (H) in the posterior part blastema (corresponding to mix of A and B populations). Finally, as expected, the posterior part blastema contains populations of gut cells, absent in the parapodia blastema (Fig. 8A).

**Figure 8.**
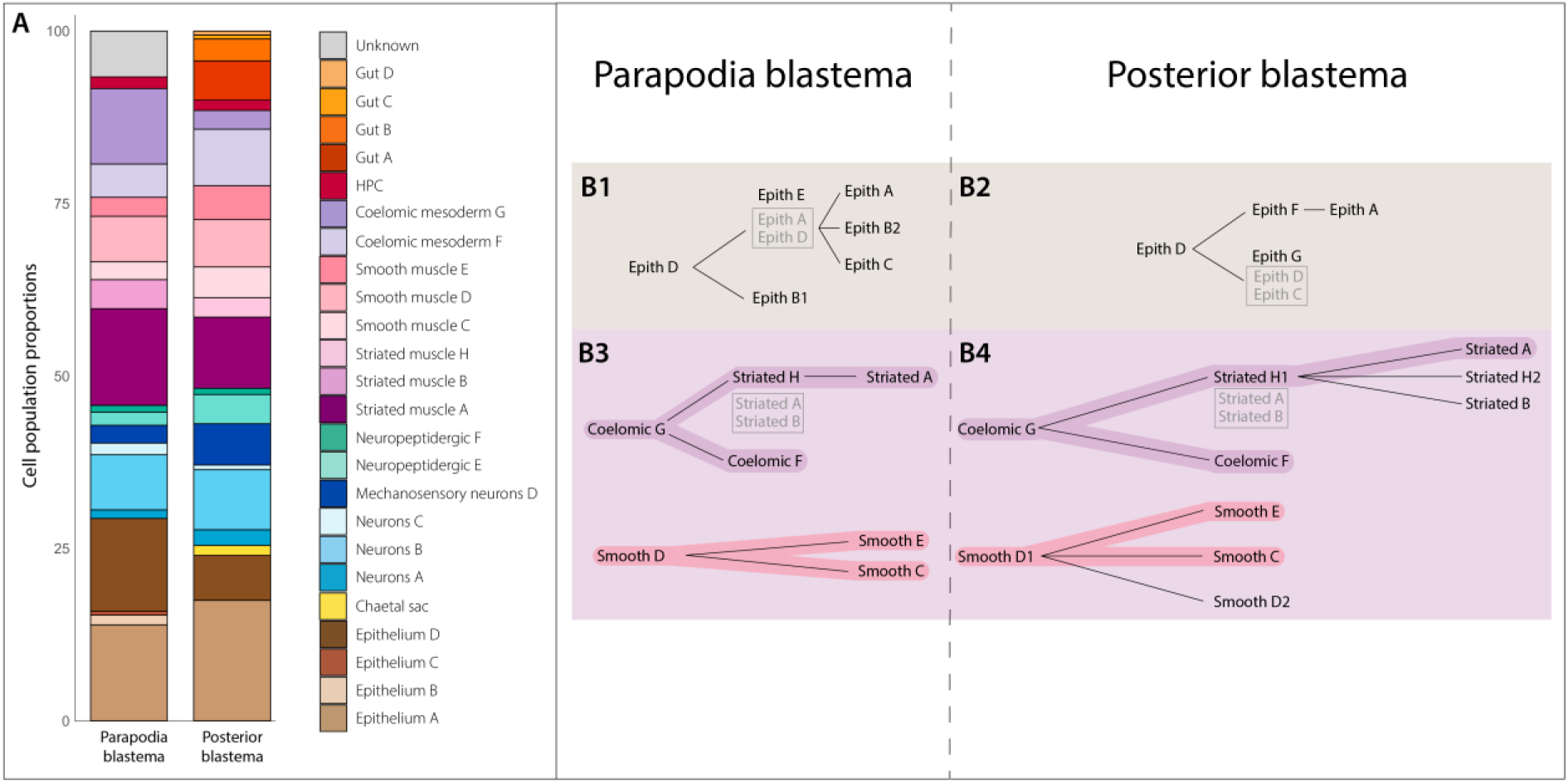
Comparison of cell populations diversity and cell differentiation trajectories between parapodial and posterior blastema. (A) Stacked plots of the main cell populations for the parapodial and posterior part blastema atlases. (B) Schematic representation of the main cell differentiation trajectories for epithelial, and mesodermal tissues, in both parapodial and posterior part blastema. Similar trajectories between the two structures are highlighted.

While the trajectories obtained for both structures are partial, as only the blastema stage is analyzed, comparison between them revealed both divergences and many commonalities (Fig. 8B). At the level of the epithelial populations, a similar putative progenitor population (Epithelium D, expressing *Sp/Btd, Flrtl* and *Evx*) was found in both the parapodia and posterior part blastema. However, the intermediate cell populations and the more differentiated epithelial cells they give rise to are different (Fig. 8B1, B2). The trajectories of mesodermal tissues appear similar at a global scale. The putative populations of broad mesodermal progenitors (Coelomic mesoderm G, expressing *Prrx*, *P2x*, and *Chd3/4/5b*) can give rise to the second population of Coelomic mesoderm (F) as well as to Striated muscle A populations, through an intermediate population of Striated muscle H. In addition, Smooth muscle population D may lead to Smooth muscles C and E (Fig. 8B3, B4). It remains to be determined whether smooth muscles are produced by coelomic mesoderm cells or are of independent origins.

Through this comparison between posterior and parapodial blastema, we determined potentially analogous progenitor cells for the main tissues and identified similar, although partial, cell differentiation trajectories. Hence these results highlight the commonalities of complex structures regeneration in *Platynereis*.

## Discussion

### Parapodia are a highly complex structure of the annelid body plan

Body appendages are major and diverse structures of bilaterian animals whose evolutionary history remains controversial (Prpic, 2019). Annelid body appendages or parapodia are biramous structures flanking the worm trunk bearing chitineous bristle-like structures named chaetae (Fig. 1B_2_, B_5_) (Grimmel et al., 2016; Paululat & Purschke, 2025). Ancestrally present in annelids (Parry et al., 2014), parapodia ensure various functions of locomotion, respiration and sensory perception (Brusca et al., 2023; Paululat & Purschke, 2025). Parapodial external and gross morphologies have been thoroughly described in a variety of annelids over the decades (Allentoft-Larsen et al., 2021; Horridge, 1962; Lawry, 1967). However, the exact organization and composition of such structure at a deeper level are only partially understood, and while descriptions of the musculature and connectome of three segmented larvae have been explored and allowed for the description of their internal composition (Jasek et al., 2022; Verasztó et al., 2025), it is unknown if this organization is conserved in juveniles. Here, we combined classical histology and microscopy approaches, tissue-specific fluorescent imaging and single cell transcriptomic (scRNAseq) to provide an understanding of juvenile parapodia at an unprecedented resolution (Fig. 1). In our cell atlas, we uncovered 20 broad cell populations and up to 45 fine-grained cell populations – encompassing the expected epithelium, nervous-system related and mesodermal tissues (Fig. 1, Supp. Fig. 4).

Parapodia epithelium comprises the epidermis covered by a collagenous cuticle and the epithelium lining the various glands (Paululat & Purschke, 2025). The two types of glands born by the parapodia (flask-shaped and tubular-cell) have been described at the morphological level, but their functions remain obscure aside for the production of different muccopolysaccharides (Daly, 1973). Here, we identified at least 12 epithelial cell populations, one of them (Epithelium 2) may be progenitor cells as it expresses members of the Germline Multipotency Program (GMP, a molecular signature of adult progenitors and stem cells (Fierro-Constaín et al., 2017; Juliano et al., 2010)), an ectodermal progenitor marker (*Sp/Btd*, (Stockinger et al., 2024)), as well as cell cycle genes (Supp. Fig. 4A-D)). Those cells may generate new epidermis during the continuous transverse growth of the juvenile parapodia. We also found a population of chaetal sacs cells, the structures producing the chaetae. However, we failed to isolate the different cell populations representing the known diversity of these structures (*i.e.* the chitin-producing chaetoblast and its associated diverse follicle cells (Gazave et al., 2017; Tilic et al., 2023; Zakrzewski, 2011)).

Parapodia are highly sensory structures. All cirri and lobes, as well as external and internal chaetae are innervated and connected to the ventral nerve chord and dorsal nerves (Fig. 1B_3_, Supp. Fig. 1B). Accordingly, we found a high number (21) of nervous system-related cell populations (Supp. Fig. 4E-H). We figured out that some of them are neuropeptidergic, photosensory and mechanosensory neurons (including the ones responsible of the larval startle response (Bezares-Calderón et al., 2018)). We also found a very small population of less differentiated neurons representing a putative neural progenitor population, and many *a priori* differentiated neuron populations that remain to be precisely identified. To this end, crossing the expression data contained in our single cell atlas to existing databases for *Platynereis* larvae such as ProSPr that contains the molecular identities of hundreds of cells from the ventral nerve chord (Vergara et al., 2017), or whole body connectomics defined in exquisite details through electron microscopy reconstructions (Verasztó et al., 2025; Williams et al., 2017) would be a powerful approach (Williams & Jékely, 2019).

Parapodia are also very mobile structures, thanks to a diversity of muscles allowing movements of each lobe as well as internal and external chaetae (Supp. Fig. 1C_1_-C_3_) (Tzetlin & Filippova, 2005). This diversity is reflected at the cellular level, as we identified up to eight muscle cell populations, half of which were striated and half of which were smooth (Supp. Fig. 4I-L). Surprisingly, based on analyses of musculature of 6 days post fertilization larvae, *Platynereis* smooth muscles were considered to be restricted around the gut (Brunet et al., 2016). Our single cell atlas challenges this view as we found a relatively important proportion of smooth muscles cells in parapodia, although they are less prominent than striated ones. Where those smooth muscles are localized remains to be determined using molecular markers. Also, given that both types of muscles are very close together, especially around the midgut, we cannot exclude the possibility that visceral muscles extend more laterally than expected and are thus collected when recovering parapodia samples. In addition, we found an interesting cell population that was previously coined as “coelomic mesoderm” (*i.e.*, “a sub-epidermal mesodermal cell type that does not include muscle”) during posterior regeneration and elongation (Fig. 1C, D) (Stockinger et al., 2024). Those cells have been proposed to participate in the regeneration of the mesodermal posterior stem cells responsible for the worm posterior growth (Gazave et al., 2013; Planques et al., 2019; Stockinger et al., 2024). In a non-amputated parapodia, those cells overexpress many GMP markers, several mesodermal progenitor markers as well as cell cycle genes (Supp. Fig. 3B). The function of such cells in parapodia is unknown, but they may participate in their transverse growth by providing new mesodermal cells.

Last, parapodia serve as gills ensuring respiration and gas exchanges through a rich network of vessels and capillaries (Song et al., 2020). In our atlas, we identified HPC or hemoglobin-producing cells (Fig. 1C, D). They are specialized cells lining the vessels, part of the mesodermal epithelium, that secret extracellular globins towards the vessel lumen (Song et al., 2020).

Altogether, our atlas recapitulates the cell type diversity expected for such a complex structure ensuring multiple functions. While we strived for a precise assignation of all the cell populations we found, many of the specific genes they overexpress are yet to be identified, and their expression pattern described. Hence, this detailed cell atlas provides a significant resource for future more specialized studies on parapodia and overall tissue composition in *Platynereis*.

### Parapodial regeneration is stereotypic and follows the main metazoan regeneration steps

Why some species are able to successfully regenerate complex structures such as appendages, while others cannot, is a central question in biology. Here, we present a detailed characterization of *Platynereis* parapodia regeneration and provide the essential elements for comparative studies of such a process in and beyond *Platynereis*.

Amputation of a parapodium leads to the removal of the two lobes bearing the sensory cirri, including the flask-shaped and tubular-cell glands, the chaetal sacs and their mobile chaetae, and induces the break of the strong internal structures providing support to parapodia, the aciculae. In 16 days, a new parapodium comprising all the aforementioned structures, while smaller to the initial one, is reformed (Fig. 2). It is well integrated in the remaining trunk tissues and ensures, at least, its locomotion function, as it is perfectly mobile and coordinated with the other parapodia. While we did not assess directly the sensory and respiratory functions, we assume that they are also restored, as the innervation of the structure and production of extracellular globins are reestablished by the end of the process (Supp. Fig. 6H_6_, I_6_). Based on morphological observations, we defined five regeneration stages and, as far as homogenous worms in size and age are concerned, the timeline of these stages appears highly reproducible (Fig. 2). This long phase of regeneration is followed by a post-regenerative growth phase of variable length lasting roughly five days, until the newly formed parapodium becomes indistinguishable from the others. This blastema-based regeneration process and its associated stages follow the three key steps of regeneration and present the hallmarks of animal regeneration (Bideau et al., 2021; Poss & Tanaka, 2024; Tiozzo & Copley, 2015).

- <u>Wound healing and regeneration initiation</u>: 2 to 3 days post-amputation (dpa, stage 1), an epithelium (named wound epithelium) covers the injury. Cell proliferation starts to increase in this epithelium and its underlying cells, while this proliferation is dispensable for regeneration to proceed further (Fig. 2B’’, I). The wound epithelium also expresses many parapodia primordia markers (*Dlx, Zic* and *Sfrp1/2/5*) and nervous-system related genes (*Ngn, Elav* and *Syt*) (Fig. 3A_2_, D_2_). In the same line, nerves project, from non-amputated tissues of the trunk, toward this epithelium (Supp. Fig. 6N_2_). This would be consistent with the hypothesis that *Platynereis* parapodial regeneration, like that of many other species, could be nerve-dependent (Boilly et al., 2017; Sinigaglia & Averof, 2019). In contrast, muscles retract while muscle markers are expressed in the underlying cells (Fig. B_2_, Supp. Fig. 6G_2_).
- <u>Mobilization of precursors cells:</u> 4 to 6 dpa, like in many annelid regeneration contexts, a blastema-like organization is formed upon parapodia amputation (Morgan, 1901; Özpolat & Bely, 2016). It corresponds to a mass of highly proliferative cells, lined by the wound epithelium, expressing broadly cell cycle and GMP markers genes (Fig. 2C”, I, Supp. Fig. 5A_2_, 8). From this stage, cell proliferation is essential for blastema growth and latter steps of morphogenesis (Supp. Fig. 5C). Parapodia blastema is not a homogenous structure. As for the non-amputated parapodia, it contains epithelium, neurons, neuropeptidergic, muscles (smooth and striated), and coelomic mesoderm cell types (Supp. Fig. 2D, 7D). Among them, several progenitor populations, more or less engaged in differentiation and specific to the epithelium, mesoderm, neurons, smooths muscles are found (Fig. 7). Surprisingly, although the external morphology of the structures does not reflect any kind of morphogenesis (Fig. 2C), many cell populations have already differentiated, particularly for the nervous system whose axonal projections extend extensively into the blastema (Supp. Fig. 6N_3_). Similarly, the muscles fibers spread into the proximal part of the blastema (Supp. Fig. 6G_3_).
- <u>Morphogenesis:</u> Even if cell differentiation starts very early, a 10-days long morphogenesis step is necessary to reform a complete parapodia. This long process allows the fine reformation of the chaetal sacs and the chaetae, then the individualization of the sensory cirri and the lobes and finally the reformation of the two types of glands (Fig. 2D-G”). Concomitantly, the whole structure is innervated, muscularized and vascularized (Fig. 3 A_4_-D_7,_ Supp. Fig. 6G_4-7_, N_4-7_). How the dorso-ventral and proximo-distal axis are reestablished during parapodial regeneration remains unknown. However, surgical experiments performed on a closely related species of *Platynereis, Nereis pelagica*, identified the ventral nerve chord (VNC) as a signaling structure establishing ventral identity, essential for the correct formation of parapodia, during posterior regeneration (Boilly et al., 2017). During parapodial regeneration, the ventral lobe differentiates slightly before the dorsal one, as evidenced by its earlier chaetae reformation (Fig. 2D, D’). However, it remains to be determined if the nerves coming from the ventral parapodial ganglia provide important cues for parapodial regeneration and dorso-ventral determination.

### Parapodial and posterior part regeneration: similarities and specificities

*Platynereis* worms, like many annelid species, harbor important regenerative capabilities (Özpolat & Bely, 2016; Schenkelaars & Gazave, 2021). These worms can regenerate two types of complex structures: their parapodia, composed of a diversity of differentiated cell types and few progenitor populations, and their posterior part comprising two populations of posterior stem cells (responsible of the worm’s continuous growth through posterior elongation), in addition to pygidial differentiated cells (Gazave et al., 2013; Planques et al., 2019). *Platynereis* constitutes a key model system where it is possible to compare the regeneration of two different structures with the aim of highlighting the elements important for this regenerative success.

For both complex structures, regeneration is successful and leads to, *a priori,* the perfect reformation of the amputated structures. Those two processes recapitulate the three key regeneration steps, are blastema-based, and are cell proliferation dependent - from the moment the blastema begins forming. However, the timing of the whole process of regeneration differs a lot. Posterior regeneration is very rapid and terminates within 5 days. The wound healing step is over at 1 dpa, the blastema is formed and grows in two more days, followed by two days of morphogenesis. Conversely, for parapodial regeneration, wound healing and blastema formation take roughly twice as long as for posterior regeneration, and the morphogenesis step is extremely long (10 days). This might be due to the complexity of a parapodia which, at least at the morphological level, appears much higher compared to the posterior part. Differences are also present for muscle and nervous system regeneration. During posterior part regeneration, the nerves grow into the blastema only from the VNC, while for the parapodia, both dorsal and ventral nerves participate. During posterior regeneration, muscles of the pygidium form *de novo*, while for parapodia regeneration, muscles fibers of the trunks extend and invade the regenerating parapodia.

The difference of morphological complexity between the two structures is partly reflected at the cellular level. When comparing blastema produced during posterior and parapodial regenerations, their cell type diversity, at a broad level, appears more or less similar (for the comparable tissues, as no gut cells are found in the parapodia blastema) (Fig. 8A). There is, however, one noticeable exception for the epithelial cell types, which are more diverse in the parapodia blastema. This may be related to the specific parapodial glands. It is worth noting that we made this comparison of cell type diversity only at this blastema stage of regeneration, which does not reflect the diversity of differentiated cells of the fully regenerated structures. It would be very interesting to generate a cell atlas of the posterior growth zone and pygidium, to compare with the one we obtained for the parapodia. In addition, we found two populations of broad ectodermal and mesodermal progenitors (Epithelium D2 and Coelomic mesoderm G, respectively), and two populations of more engaged neural and smooth muscles that are transcriptomically similar between the two blastemas. They overexpress key progenitor markers, members of the GMP signature and cell cycle genes (Supp. Fig. 4, 9). Putative common cell differentiation trajectories were also identified for the neurons, the coelomic mesoderm and smooth muscles (Fig. 8B). However, they differ for the epithelial cells. We also lack information about the cell trajectories leading to the formation of the neuropeptidergic and two gut cell populations. As these cell types appear already well differentiated at the blastema stage, the cell trajectories we identified could not attribute these cell populations to any given trajectory already present. In addition, if smooth muscles originate from coelomic mesoderm, as the striated ones, or have a different origin remain to be determined.

In a diversity of annelid species, cell dedifferentiation was often proposed as the major source of blastema cells (Kostyuchenko & Kozin, 2021; Planques et al., 2019; Stockinger et al., 2024). *Platynereis* posterior regeneration relies on ectodermal, mesodermal and endodermal lineage-restricted progenitors probably produced through dedifferentiation of wound-adjacent cells, located in the segment abutting the amputation plan (Bideau et al., 2024; Planques et al., 2019; Stockinger et al., 2024). The presence of highly similar ectodermal and mesodermal progenitors in the parapodial blastema suggests that parapodial regeneration could also rely on dedifferentiation to produce blastema cells. However, early cell trajectories need to be established to resolve this question.

In this study, we explored the mechanisms of complex structure regeneration in a single well-suited species, the annelid *Platynereis dumerilii*. Through an original comparative approach between parapodia and posterior regenerations, we identified the major cell types participating in successful regeneration events in this species. Our data provided a fresh perspective on fundamental regenerative processes which in term will help unravel regeneration success in animals.

## Material and methods

### *Platynereis dumerilii* breeding culture and amputation procedures

*P. dumerilii* juvenile worms were obtained from a husbandry established in the Institut Jacques Monod and Saint-Petersburg University, Russia (for precise breeding conditions, see (Dorresteijn et al., 1993; Vervoort & Gazave, 2022)). For parapodia samples, worms used were 4-5 months old and 40-60 segments long. For the individual segments shown in transversal view, the 9^th^ to 11^th^ posteriormost segments were individually collected from worms anesthetized (using a 1:1 solution of MgCl_2_ 7.5% and sea water) with a micro knife (Sharpoint™). For homeostatic parapodia samples, the 9^th^ to 11^th^ posteriormost segments were collected from. For parapodia regeneration samples, amputations were performed on the 10^th^ posteriormost segment of curled worms, to remove as much parapodial tissue as possible without carving into the trunk of the animals, as is depicted by the dotted grey lines in Figure 2. Posterior amputations were done according to the procedure detailed by (Planques et al., 2019; Vervoort & Gazave, 2022) 3-4 month old worms with 30-40 segments. After amputation, worms were left to recover in fresh natural sea water and fed three times per week.

### Scanning electron microscopy (SEM)

Juvenile worms (in homeostasis or at defined stages of parapodial regeneration) were collected, relaxed in 7,5% MgCl_2_•6H_2_O, and fixed in 2.5% Glutaraldehyde prepared with 0.05M Sorensen’s phosphate buffer (pH=7,2) and 0.3M NaCl. Then they were washed in 0.05M Phosphate buffer with 0.3M NaCl dehydrated in ascending alcohol series, immersed in acetone, critical point dried, coated with platinum, and examined under FEI Quanta 250 scanning electron microscope. (Starunov et al., 2015).

### Semithin section and 3D reconstructions

For 3D-reconstructions, samples were fixed in in 2.5% glutaraldehyde buffered in 0.05 M Sorensen’s phosphate buffer (pH=7,2) and 0.3 M NaCl for 2h, rinsed 3 times in the same buffer, and postfixed in 1% OsO4 in the same buffer for 1 h. The specimens were dehydrated in ascending acetone series, transferred in propylene oxide and embedded in Epon-Araldite resin (EMS, 13940). A series of semithin (700 nm) sections were cutusing either a Reichert Ultracut E or Leica Ultracut UC6 and a Diatome HistoJumbo diamond knife (Blumer et al., 2002) stained with methylene blue and basic fuchsine (D’Amico, 2005) and imaged using a Leica DM2500 microscope. The images were aligned with IMOD and ImodAlign tool. The 3D reconstructions and videos were made with Fiji TrackEM2 plugin.

### Scoring

For various experiments, worms were observed under a dissecting microscope at different time points after amputation, and each was attributed a score based on the regeneration stage exhibited. If worms exhibited intermediate morphologies between stages, they were scored as halved numbers *(i.e.*, 0.5,1.5, …).

### Biological material fixation (whole mount)

For most experiments performed (*i.e., in situ* hybridizations, antibody staining and EdU assays), samples were collected and fixed in 4% paraformaldehyde (PFA) diluted in 1X PBS Tween20 0.1% (PBT) for 2 h at room temperature, then washed in 1X PBT and gradually transferred to 100% methanol, at which point they were stored at −20°C. For phalloidin labeling, samples were similarly fixed but not put on methanol, instead they were kept in 1X PBT at 4°C for up to 4 days before labelling.

### Whole mount *in situ* hybridization (WMISH), antibody staining and phalloidin labeling

Probe synthesis and colorimetric NBT/BCIP WMISH and immunolabelling were performed as previously described (Demilly et al., 2013; Gazave et al., 2013; Vervoort & Gazave, 2022). Following rehydration, samples were treated with 40 µg/ml proteinase K in PBT for 10min, 2mg/ml glycine PBT for 1min, 4% PFA PBT for 20min and washed in PBT prior to hybridization or labelling. For hybridization, samples were incubated in hybridization buffer at 65°C (1.5 h), then incubated overnight with digoxigenin (DIG)-labelled probes at 65°C. DIG was then bound to specific antibodies bearing alkaline phosphatase and the tissues were stained by cleavage of NBT (Nitro Blue Tetrazolium chloride) by alkaline phosphatase, catalysed by BCIP (5-bromo-4-chloro-3-indolyl phosphate). Samples were then mounted in glycerol for imaging under a bright field microscope.

Neurite labelling was done as previously described (Demilly et al., 2013; Gazave et al., 2013), using mouse anti-acetylated tubulin (Sigma 1:500) antibodies and fluorescent secondary anti-mouse IgG Alexa Fluor 488 conjugate (Invitrogen, 1:500). For phalloidin labelling, samples were incubated in phalloidin-Alexa 555 (Molecular Probes, 1:100) overnight at 4°C. Next, samples were nuclei counter-stained with Hoechst 0.1% overnight at 4°C and mounted in glycerol/DABCO (2.5mg/ml DABCO in glycerol) for confocal imaging.

### EdU cell proliferation assay

Proliferating cells were labelled by incubating worms with 5 µM of the thymidine analog 5-ethynyl-2′-deoxyuridine (EdU), for 1h in sea water prior to fixation. Fixed samples were subsequently fluorescently labelled with the Click-it EdU Imaging Kit (555 nm, ThermoFisher) and nuclei were counter-stained with Hoechst 0.1% overnight at 4°C and mounted in glycerol/DABCO (2.5mg/ml DABCO in glycerol) for confocal imaging, as previously described in (Planques et al., 2019).

### Cell proliferation inhibitor treatments

Cell proliferation during regeneration was blocked using a well characterized inhibitor, hydroxyurea (HU, Sigma H8627) efficient in *Platynereis* (Planques et al., 2019). HU was dissolved in sea water (10mM). Worms were incubated in 3mL of HU solution in 6-well plates and the HU solution was changed every 24h to maintain its activity for the duration of the experiment.

### EdU+ cell counting

EdU cell counts were performed using the Imaris software by BitPlane (version 10.0) following the automatic cell counting procedure defined in (Bideau et al., 2024). Briefly, Hoechst-stained nuclei were used to define a region of interest (ROI) corresponding to the regenerative or homeostatic part. Then the spots inside the ROI were sorted along the EdU fluorescent signals with a filter ‘Intensity Mean’ to discriminate true positive nuclei from background.

### Statistical analysis

All statistical tests and graphic representations were performed using the Prism 9 software (GraphPad). For HU treatments and EdU+ proportions, multiple Mann -Whitney tests were performed to compare each experimental condition to the control and p-values were corrected using the Bonferroni-Dunn method.

### ACME cell dissociation

#### Samples collection

For homeostatic samples, parapodia (n=21 000) along most of the body of the animals were dissected using a micro knife. For blastema samples, posterior part (n= 580) and parapodia blastemas (n= 800) were similarly collected at 3- and 5-days post-amputation (dpa), respectively. The ACME solution (García-Castro et al., 2021) was prepared for homeostatic and blastema samples using a ratio of RNase/DNase-free water, methanol, glacial acetic acid and glycerol of 13:3:2:2 or 11:3:4:2, respectively. During collection, samples were kept on a cold ACME solution supplemented with 400U/mL of recombinant RNAse inhibitor (Promega, N2511) and 75% N-acetyl-L-cysteine (Sigma Aldrich, A7250). Afterwards, the solution was replaced by a room temperature ACME solution containing 200U/mL of recombinant RNAse inhibitor.

#### Cell dissociation

To dissociate the samples, they were shaken at 1200 rpm, 20°C for 1 hour using an Eppendorf Thermomixer and vigorously pipetted, to generate bubbles, every 10 minutes (using a P1000 pipette). After dissociation, cell suspensions were filtered through a 30 μm CellTrics filter. They were then washed twice with a cold PBS 1X BSA 1% + 200U/mL RNase inhibitor solution and using a swinging bucket centrifuge before resuspension in a PBS 1X BSA 1% + 200U/mL RNase inhibitor and DMSO (1:10) solution and storage at -70°C.

#### Flow cytometry

To eliminate cell debris, cell suspensions were sorted by FACS. To this end, a FACS buffer containing 5% BSA (Sigma, 10735078001) and 5% HePes (1M, Thermo Fisher, 15630106) in HBSS 1X (Thermo Fisher, 14025-092) and filtered using a 0.22 µm Millex™-GS seringe filter (Sigma Aldrich, SLGS02510) was used. Cells were thawed on ice and washed two times in cold FACS buffer with 200U/mL of recombinant RNAse inhibitor, again using a swinging bucket centrifuge, before being resuspended in 500 μL FACS buffer + 200U/mL of recombinant RNAse inhibitor with 1 μL of DRAQ5 (to label nuclei, Thermo Fisher, 62251) and 0.5 μL of Concanavalin-A (Con-A) conjugated with Alexa Fluor 488 (to label cytoplasm, Thermo Fisher, C11252) and incubated on ice, in the dark for 20 min. The stained cell solution was then sorted by flow cytometry using a BD FACSAria Fusion machine using the 4-Way Purity mode, resuspended in cold 4% BSA in PBS 1X + 10 μL of RNAse inhibitor, and kept on ice until library preparation.

Post FACS sorting, 111 691 cells were collected from dissociated non-amputated parapodia, 129 000 cells from posterior blastemas, and 43 000 cells for parapodia blastemas.

### Single-cell library preparation and sequencing

Single-cell RNA-seq, library preparation and Illumina sequencing were performed at the Ecole normale supérieure GenomiqueENS core facility (Paris, France). Cellular suspension (35 000 cells) was loaded on 10x Chromium instrument to generate 11 299 single-cell Gel Beads-in-emulsion (GEMs) for non-amputated parapodia, 7 882 single-cell GEMs for posterior blastemas, and 3 877 single-cell GEMs for parapodia blastemas, using the manufacter’s instructions (Chromium Next GEM Single Cell 3′ Reagents Kit v.3.1 (Dual Index)). All libraries were sequenced on a NextSeq 2000 device (Illumina) with 28 cycles of read 1, 10 cycles of i7 index, 10 cycles of i5 index and 91 cycles of read 2. Sequencing data were processed using the Cell Ranger (version 6.1.2). Count pipeline and reads were mapped on the *Platynereis* genome version 1 (Assembly Genbank ID: GCA_026936325.1, using its associated gtf annotation file v021 (Mutemi et al., 2025)).

### Single-cell RNA-seq analysis

Subsequent analysis were performed using R version 4.3.2. Count matrices were loaded into R and empty droplets were removed using the emptyDrops function of the DropletUtils package. For our blastema samples a lower threshold of 200 UMI per droplet was set. Doublets were identified using the scDblFinder package (Germain et al., 2022) and removed. Further analysis, heatmaps, UMAPs, and gene expression Dotplots were done using Seurat version 5.1.0 (Hao et al., 2024). Outlier cells were identified based on manual inspection of plots representing the number of genes, transcripts, or mitochondrial genes expressed per cell. Quality metrics, plots and cutoff thresholds for all samples are available in the project’s Github. The correlation between the normalized gene expression and three metadata variables (number of genes, transcripts and mitochondrial genes per cell) was calculated and plotted to verify that none of these variables bias the clustering.

Datasets were normalized (using LogNormalize) and variable genes were identified (using FindVariableFeatures). Next, datasets were scaled, and principal component analysis (PCA) was performed. For clustering the number of principal components (PCs) was chosen based on how many were needed to recapitulate 75% of the standard deviation of the dataset. Additionally, graphs produced by the Clustree package (Zappia & Oshlack, 2018) were inspected and used to choose a clustering resolution that better reflects known cell types.

After an initial clustering for homeostatic and blastema parapodia samples, a cluster was identified and thought to be contamination from the gut based on high expression levels of *HNF4*, a transcription factor expressed in gut cells (Achim et al., 2018). For both datasets, the cluster in question was removed before clustering parameters were recalculated. It represents 480 and 62 cells for non-amputated and blastema parapodia samples, respectively.

For posterior blastemas, we took advantage of similar datasets available from the literature (Stockinger et al., 2024) and integrated them with ours to obtain a larger one, allowing a deeper resolution. Integration of the three datasets was done using the normalized data and following the anchoring approach provided by Seurat v 5.2.0. After analyzing the bias plot, the number of PCs and resolution were chosen as previously described.

### Subclustering

To assess the cell populations of atlases at higher resolution, subclustering analysis was performed on epithelial, nervous system, or mesodermal cell populations. The entire clustered homeostatic or blastemal datasets were subsetted into the aforementioned categories before clustering parameters were recalculated for each subset as previously described.

### Cluster annotation

For all datasets, differentially expressed genes (DEG) were identified (using FindAllmarkers) through a Wilcoxon Rank Sum test. To identify overexpressed genes, a cutoff was implemented to only obtain markers with a log2 fold change > 0 and adjusted p_value < 0.05. Among those genes, known markers from the literature were selected and plotted (using Dotplot). The resulting plots were used to annotate the different clusters. Additionally, the Seurat v 5.2.0 “mapping and annotating query datasets” vignette was used to inform parapodia blastema cell population annotations based on the NA parapodia annotations, and posterior blastema cell population annotations based on the parapodia blastema annotations. For the posterior blastema dataset, the initial annotation was refined by identifying and examining the expression of newly defined, specific markers for each population in the parapodia blastema dataset.

### Defining progenitor gene modules

We used the AddModuleScore Seurat function to create a gene module comprised of known GMP signature genes, ectodermal, mesodermal and neural progenitors, and cell cycle genes. The individual cell scores where then visualized using the FeaturePlot Seurat function.

### CytoTRACE analysis

We ranked cells based on their developmental potential using the CytoTRACE package (v.0.0.3) (Gulati et al., 2020), which leverages the diversity of genes expressed per cell to predict the differentiation state of cells. CytoTRACE scores were calculated for each subclustered dataset.

### Pseudotime analysis

To determine and visualize putative lineage trajectories, the different cell subclustered populations were analyzed in pseudotime. For the different subclusters, the differentiation trajectories were predicted following the standard vignette for the Slingshot 2.10.0 package. The starting clusters were determined based on their CytoTRACE score and expression of expected marker genes (GMP signature, ectodermal and mesodermal progenitor markers, cell cycle genes).

### Image acquisition and treatments

Bright-field images of colorimetric WMISH samples were acquired with a Leica CTR 5000 microscope. Fluorescent confocal images of whole samples were acquired with either a Zeiss LSM780 or LSM980 confocal microscopes. Image processing (contrast and brightness, z-projection, and auto-blend layers) was performed using FIJI and Abode Photoshop. Figures were assembled with Adobe Illustrator.

## Data availability

All data are available in the main text and the supplementary material, scripts are available at the XXX Github repository and data deposited at the ArrayExpress repository under the XXX accession number.

## Acknowledgments

We are grateful to all past and present members of the Gazave lab for their support and suggestions on this study. We thank in particular Ombeline Lamer for generating preliminary data on the project during her internship, Dr. Yves Clément for his advices on statistical analysis and Dr. Pierre Kerner for his advices and support on the project. We are also highly grateful to Dr. Felix Simon and Dr. Konstantina Filippopoulou for their help regarding the single cell analysis. We warmly thank Dr. Nikolaos Papadopoulos (Wien University) for providing several useful gene annotations. We thank the Institute Jacques Monod animal facility staff for their help with the *Platynereis* culture. We acknowledge the ImagoSeine core facility of Institut Jacques Monod, member of France-BioImaging (ANR-10-INBS-04) and IBiSA, with the support of Labex “Who Am I”, Inserm Plan Cancer, Region Ile-de-France and Fondation Bettencourt Schueller, and in particular Nicolas Valentin for his help regarding the cell sorting

## Funding

Work in our team is supported by funding from: Labex “Who Am I” laboratory of excellence (No. ANR-11-LABX-0071) funded by the French Government through its “Investments for the Future” program operated by the Agence Nationale de la Recherche under grant No. ANR-11-IDEX-0005-01, Agence Nationale de la Recherche «STEM» (ANR-19-CE27-0027-01)), Centre National de la Recherche Scientifique (CNRS), INSB (Grant Diversity of Biological Mechanisms), Université Paris Cité, Association pour la Recherche sur le Cancer (grant PJA 20191209482), and comité départemental de Paris de la Ligue Nationale Contre le Cancer (grant RS20/75-20). ZVR has obtained a PhD fellowship from the BioSPC doctoral school, and her fourth year of PhD is supported by the EUR GENE graduate school. She also received the 2025 Young Researchers Prize from the Fondation des Treilles. The GenomiqueENS core facility was supported by the France Génomique national infrastructure, funded as part of the “Investissements d’Avenir” program managed by the Agence Nationale de la Recherche (contract ANR-10-INBS-0009).

## Supplementary figures

**Figure S1.**
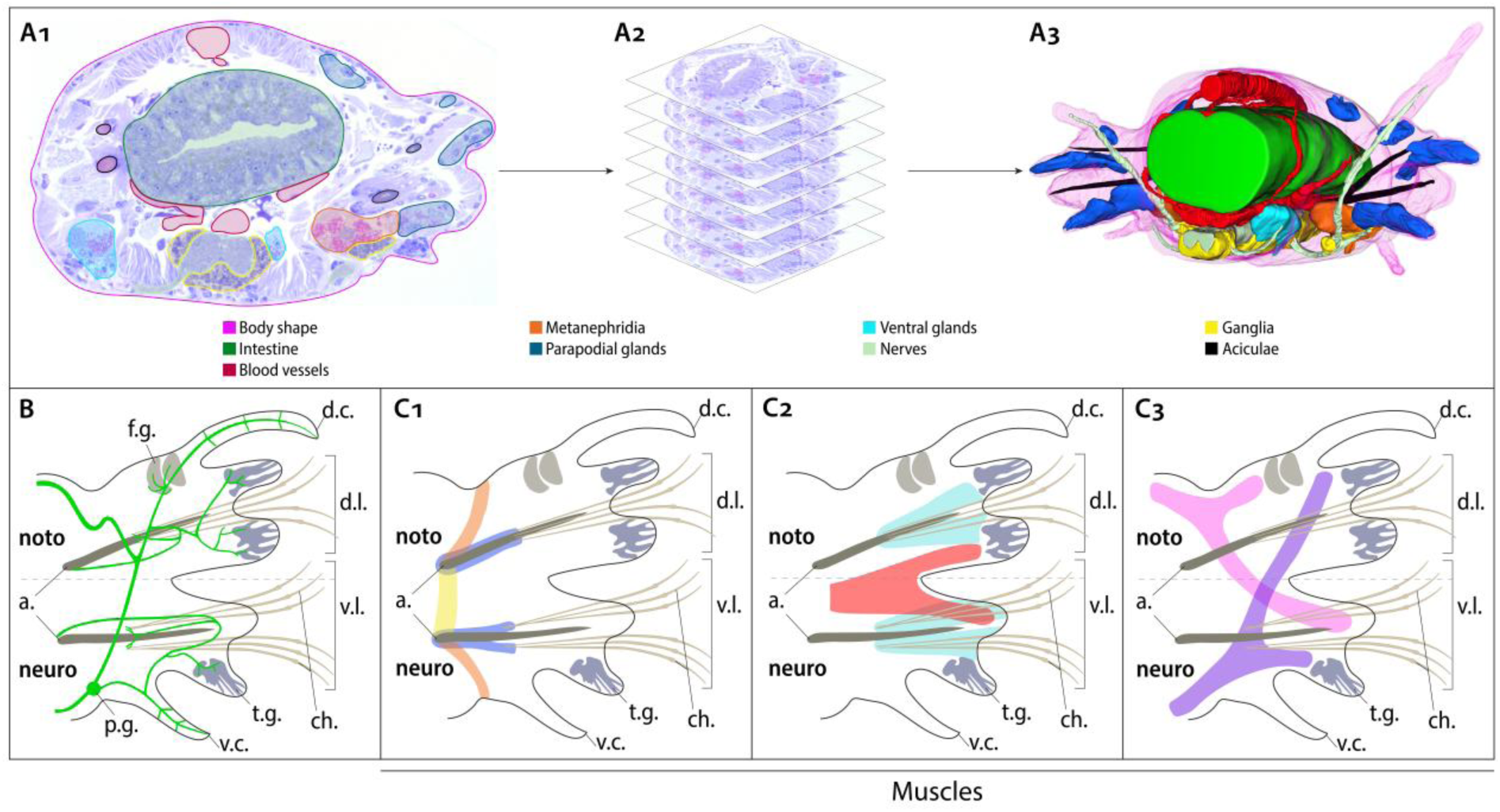
Identification of tissues/ structures and 3D reconstruction of a parapodium from semi-thin sections and fluorescent labelling. (A1-A3) Procedure for generating the 3D reconstruction. (A1) Semi-thin transversal sections are used to manually identify and outline the different tissues present. Pink, body shape; green, intestine; red, blood vessels; orange, metanephridia; dark blue, parapodial glands; light blue, ventral glands, light green, nerves; yellow, ganglia; black, aciculae. (A2) This identification of tissues is done in all images of a stack of parapodial sections and used to render a 3D reconstruction of a worm segment (A3). (B) Schematic organization of the parapodial nerves determined thanks to neurite labelling. p.g., parapodial ganglion. (C1-C3) Schematic organization of the parapodial muscles determined thanks to an actin fiber labelling. Orange: acicular muscle. Yellow: inter-acicular muscles. Dark blue: chaetal sac retractor muscles. Red: inter-acicular to lobe muscles. Light blue: chaetal protractor muscles. Pink: dorsal muscles. Purple: ventral muscles. noto. notopodium; neuro. neuropodium; a. acicula; f.g. flask-shaped gland, t.g. tubular-cell gland; d.l., dorsal lobe; d.c. dorsal cirri; v.l., ventral lobe; v.c. ventral cirri; ch. chaeta.

**Figure S2.**
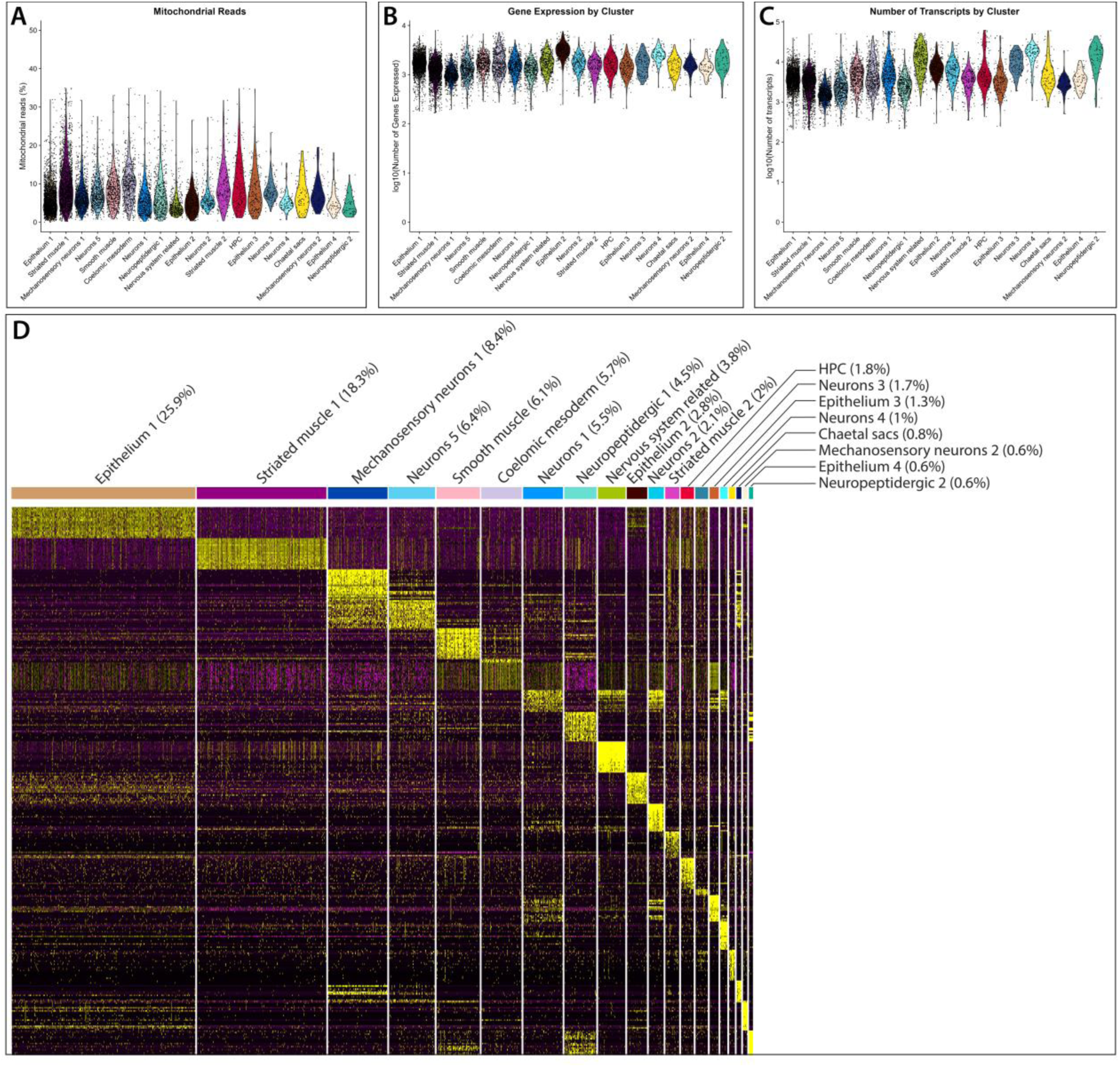
Parapodia scRNA-seq quality control metrics and gene expression across different cell populations. Percentage of mitochondrial reads (A), number of genes (B), and transcripts (C) across the identified cell populations. (D) Heatmap depicting the top 20 genes expressed per cell type for the whole atlas.

**Figure S3.**
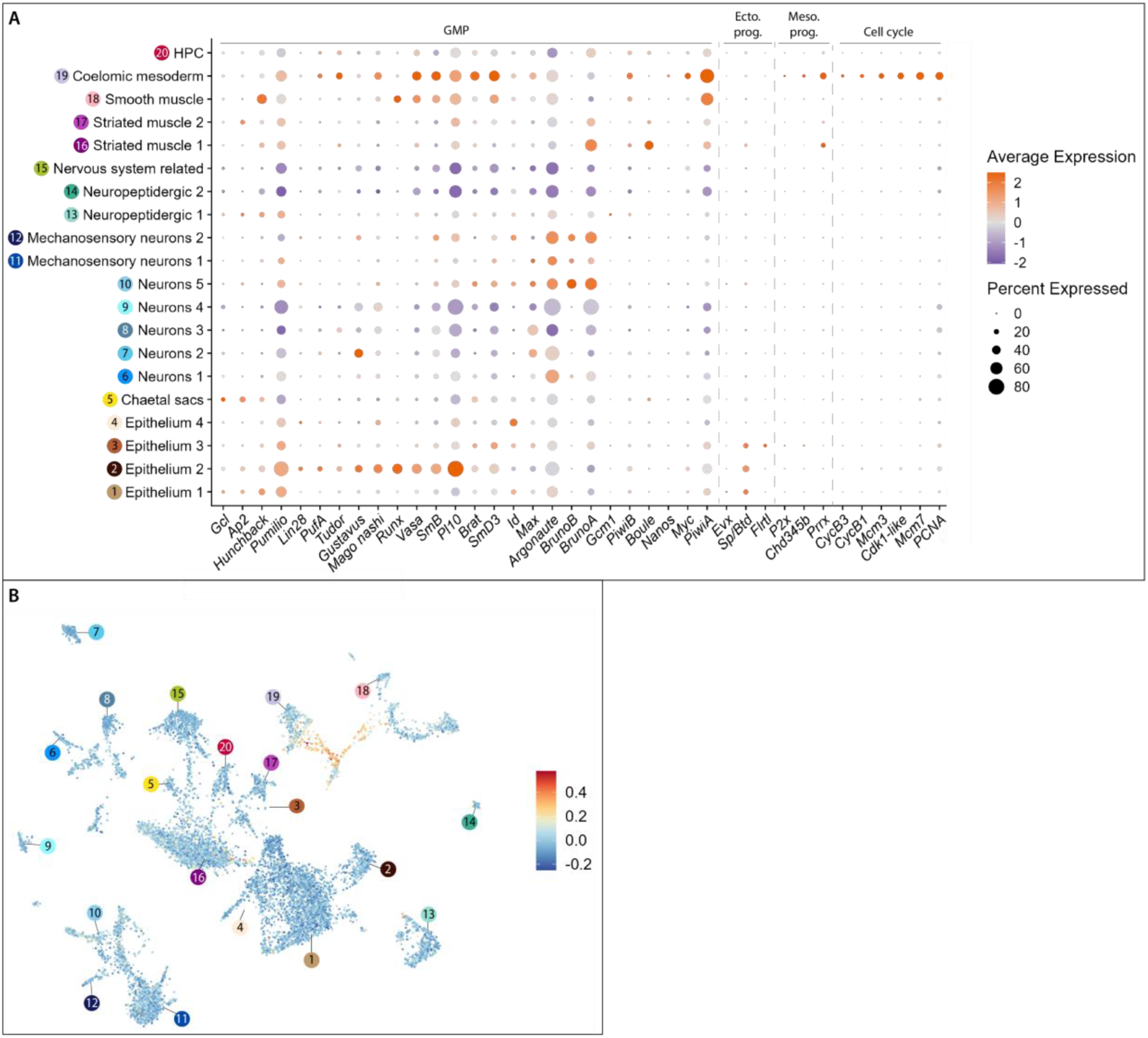
Identification of putative progenitors in the homeostatic parapodia dataset. (A) UMAP visualization of CytoTRACE values calculated for the entire dataset. High values (red) indicate less differentiated cell types. (B) Dotplot of genes belonging to the GMP signature, ectodermal and mesodermal progenitors, and cell cycle genes per cluster.

**Figure S4.**
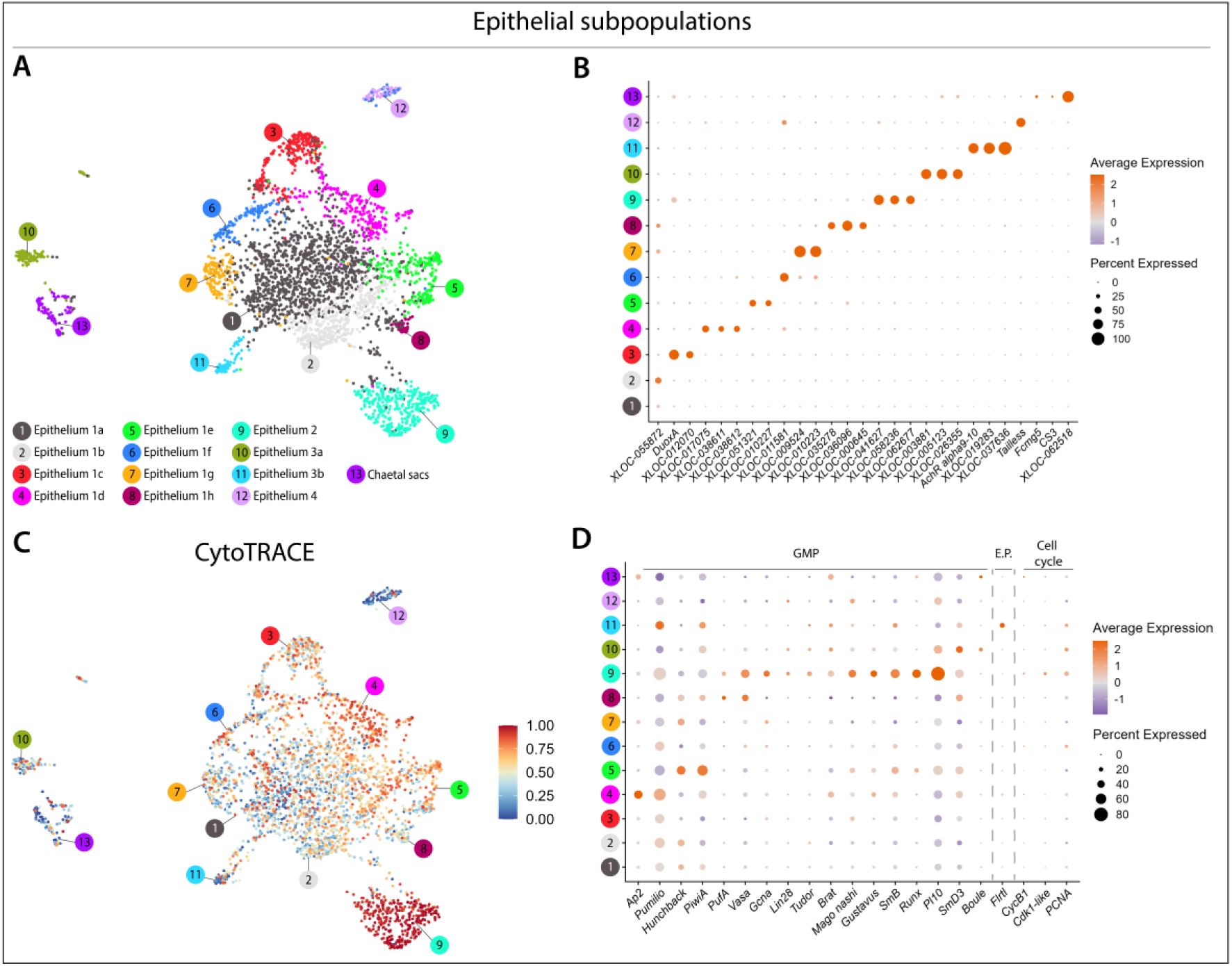
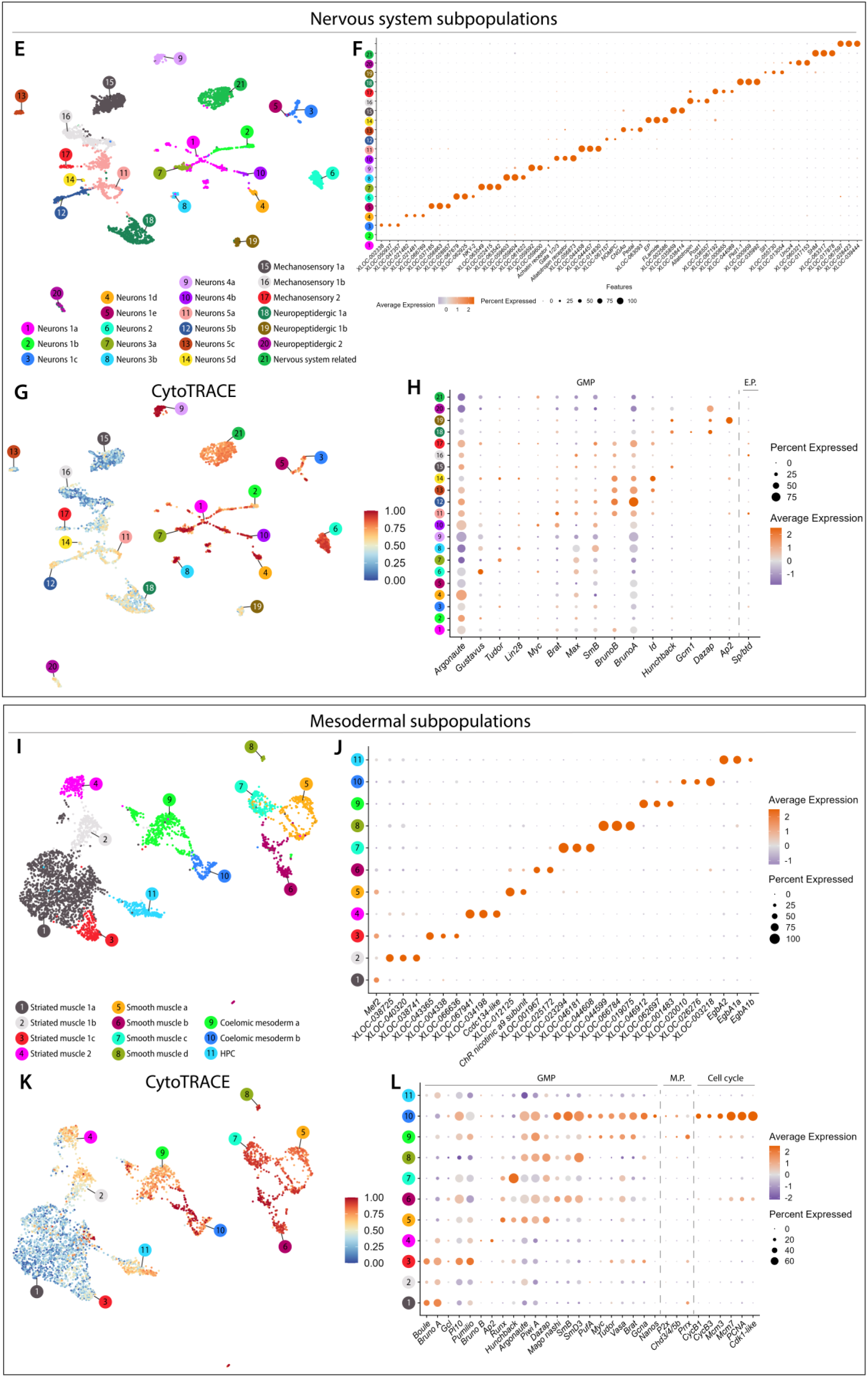
Identification of different cell subpopulations in the homeostatic parapodia dataset. (A) Subclustering of the broad epithelial cell types revealed 13 different subpopulations. (B) Dotplot of known and newly identified upregulated gene markers for each subpopulation. (C) UMAP visualization of CytoTRACE values. (D) Dotplot of upregulated genes belonging to the GMP signature, ectodermal progenitors, and cell cycle genes. (E) Subclustering of the broad nervous system subtypes revealed 21 different subpopulations. (F) Dotplot of known and newly identified upregulated gene markers for each subpopulation. (G) UMAP visualization of CytoTRACE values. (H) Dotplot of upregulated genes belonging to the GMP signature and cell cycle genes. (I) Subclustering of the broad mesodermal subtypes revealed 11 different subpopulations. (J) Dotplot of known and newly identified upregulated gene markers for each subpopulation. (K) UMAP visualization of CytoTRACE values. (L) Dotplot of upregulated genes belonging to the GMP signature, mesodermal progenitors, and cell cycle genes. E.P, ectodermal progenitors; M.P, mesodermal progenitors.

**Figure S5.**
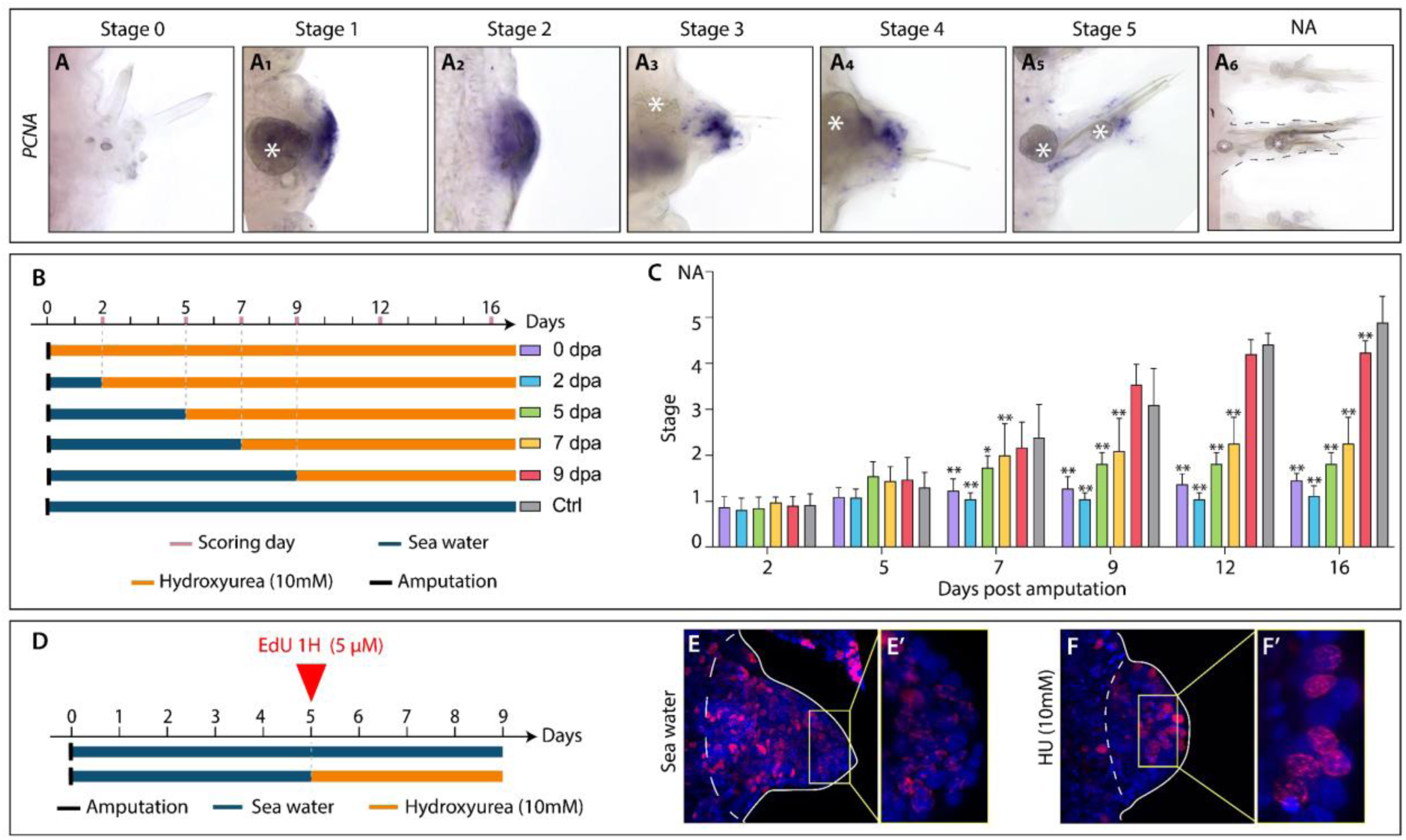
Cell proliferation during parapodia regeneration. (A-A6) WMISH for cell cycle gene PCNA shown for all six stages of regeneration and in a non-amputated parapodium. (B) A schematic representation of HU treatments. (C) Barplot recapitulating the stages reached by the worms in the different conditions (n≥11 per HU condition, n=22 for control condition). *P<.05 **P<0.001 (Mann-Whitney test with Bonferroni-Dunn correction). (D) Schematic representation of the experimental design used to confirm hydroxyurea (HU) anti-proliferative effect. 5 dpa worms were incubated one hour with 5µM EdU and chased in normal seawater (control; C-C’) or seawater with 10mM HU for four days (until 9 dpa) before observation. (E-F’) EdU labeling of control (E) and HU-treated worms (F). Zoom of some nuclei in control (E’) and HU-treated worms (F’). Dpa: days post amputation; HU: hydroxyurea; NA: non-amputated; Ctrl: control.

**Figure S6.**
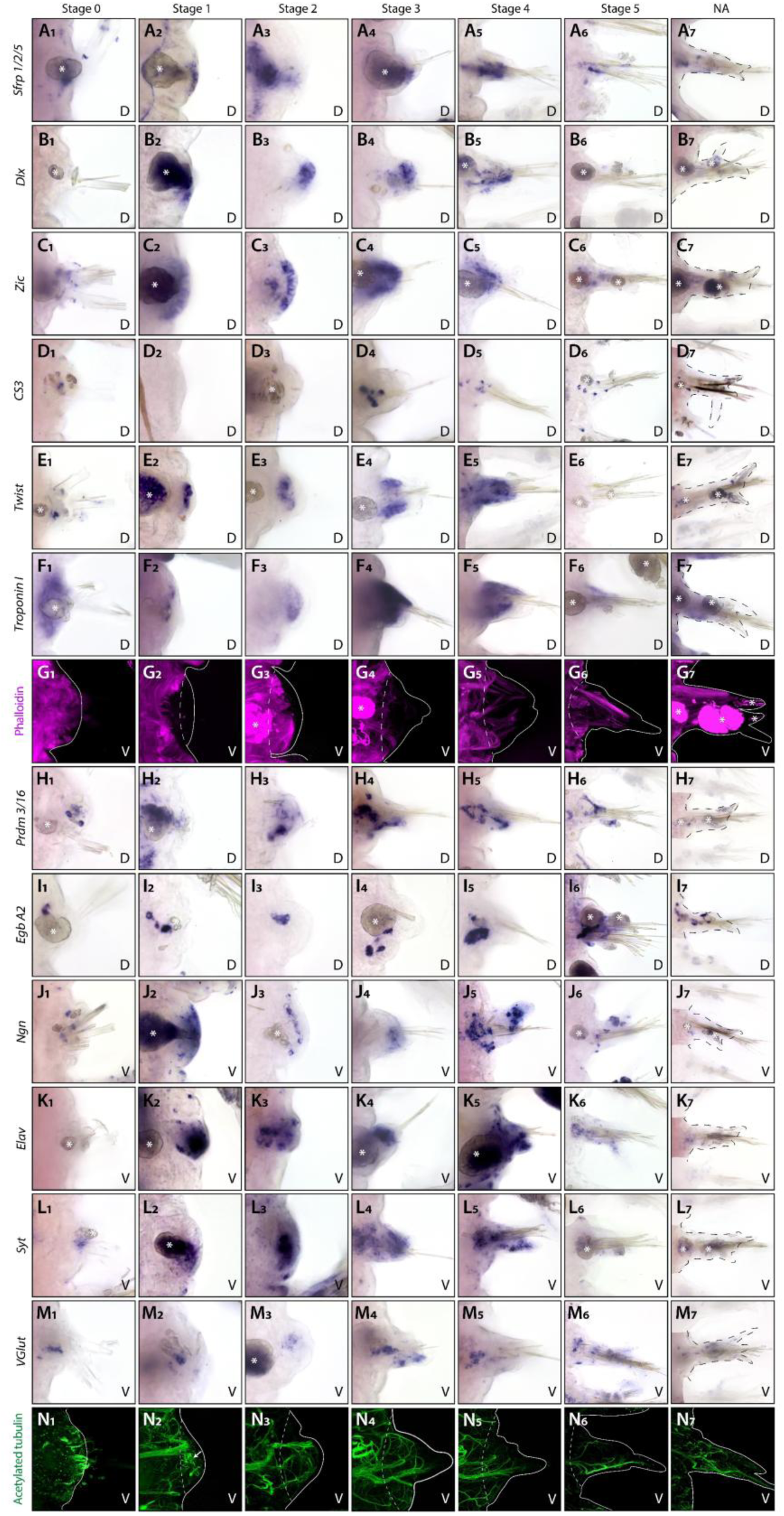
Tissue patterning and differentiation during parapodial regeneration. (A1-G7; I1-O7) WMISH of the indicated genes are shown for all six regeneration stages and for NA parapodia. Black dashed line, parapodia outlines. (H1-H7) Phalloidin labeling of muscle fibers. (P1-P7) Immunostaining of acetylated tubulin labeling the regenerating peripheral nerves. V, ventral view; D, dorsal view. White asterisks, non-specific staining of the parapodial glands; white lines, parapodia outlines; white dashed lines, amputation site; white arrow, disorganized nerves. NA: non-amputated.

**Figure S7.**
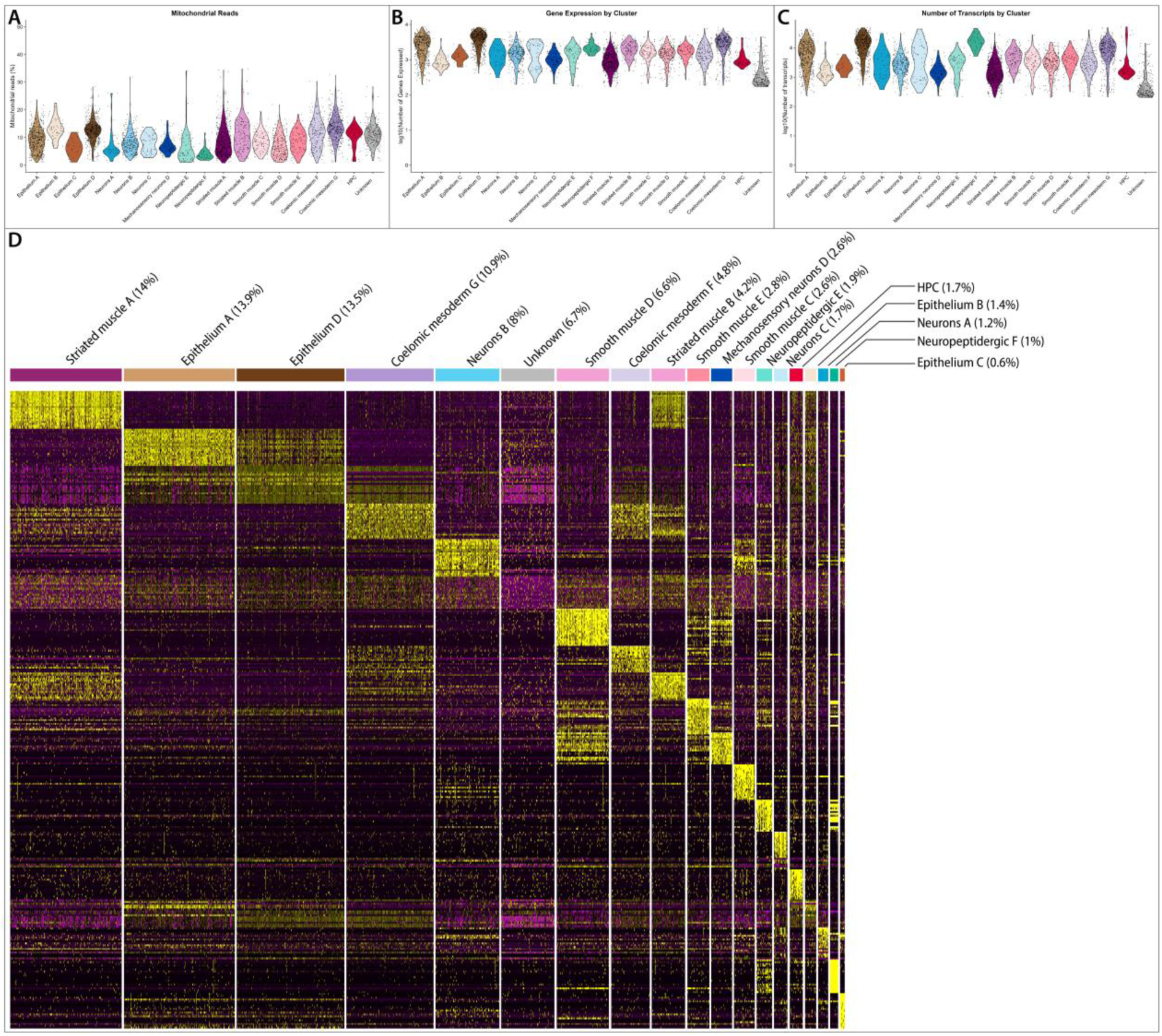
Parapodia blastema scRNA-seq quality control metrics and gene expression across different cell populations. Percentage of mitochondrial reads (A), number of genes (B), and transcripts (C) across the identified cell populations. (D) Heatmap depicting the top 20 genes expressed per cell population.

**Figure S8.**
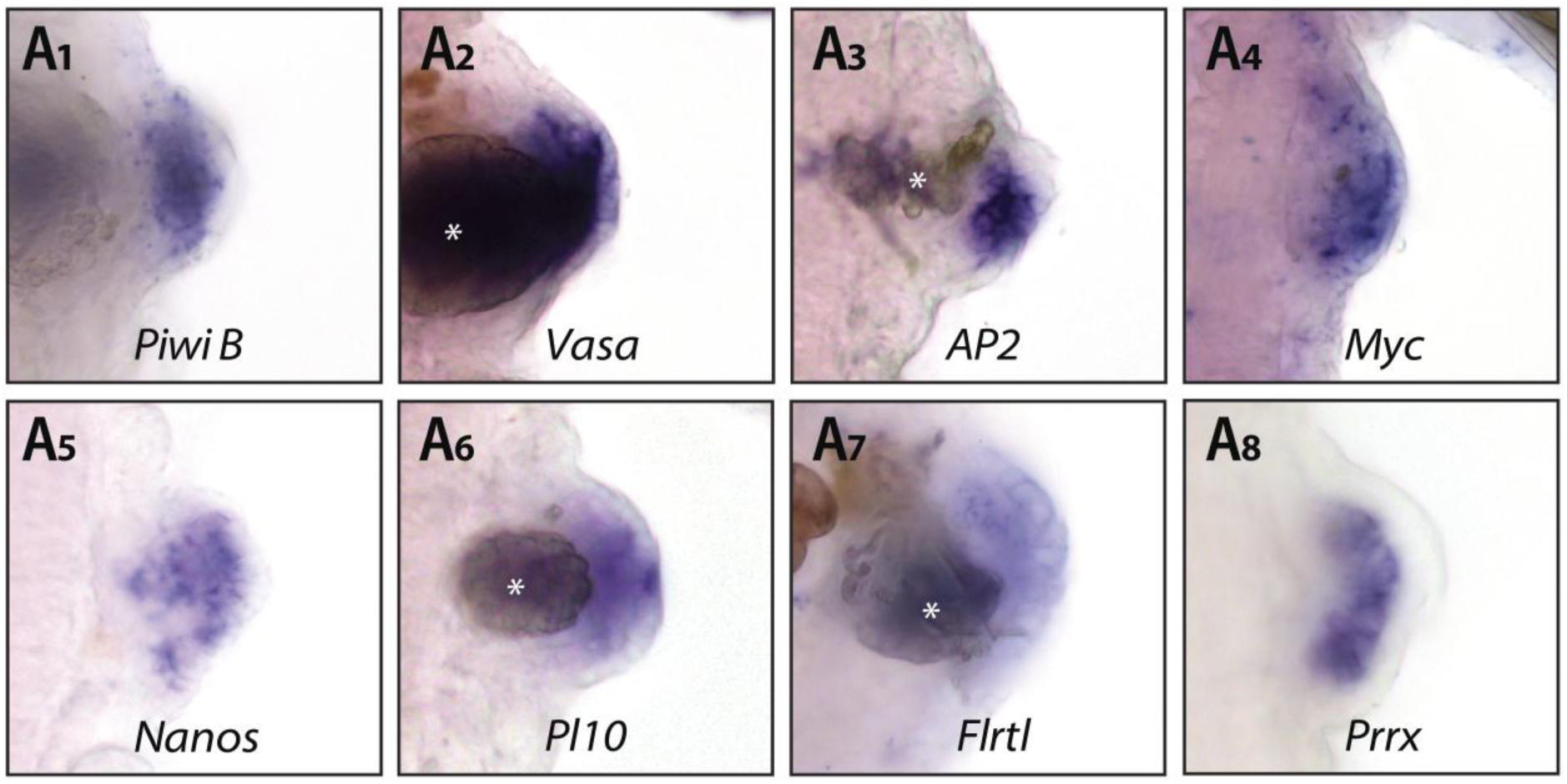
Expression of GMP markers and progenitors in the parapodia blastema. WMISH of the indicated genes are shown for the parapodia blastema (stage 2). All panels are dorsal views. White asterisks indicate the glands.

**Figure S9.**
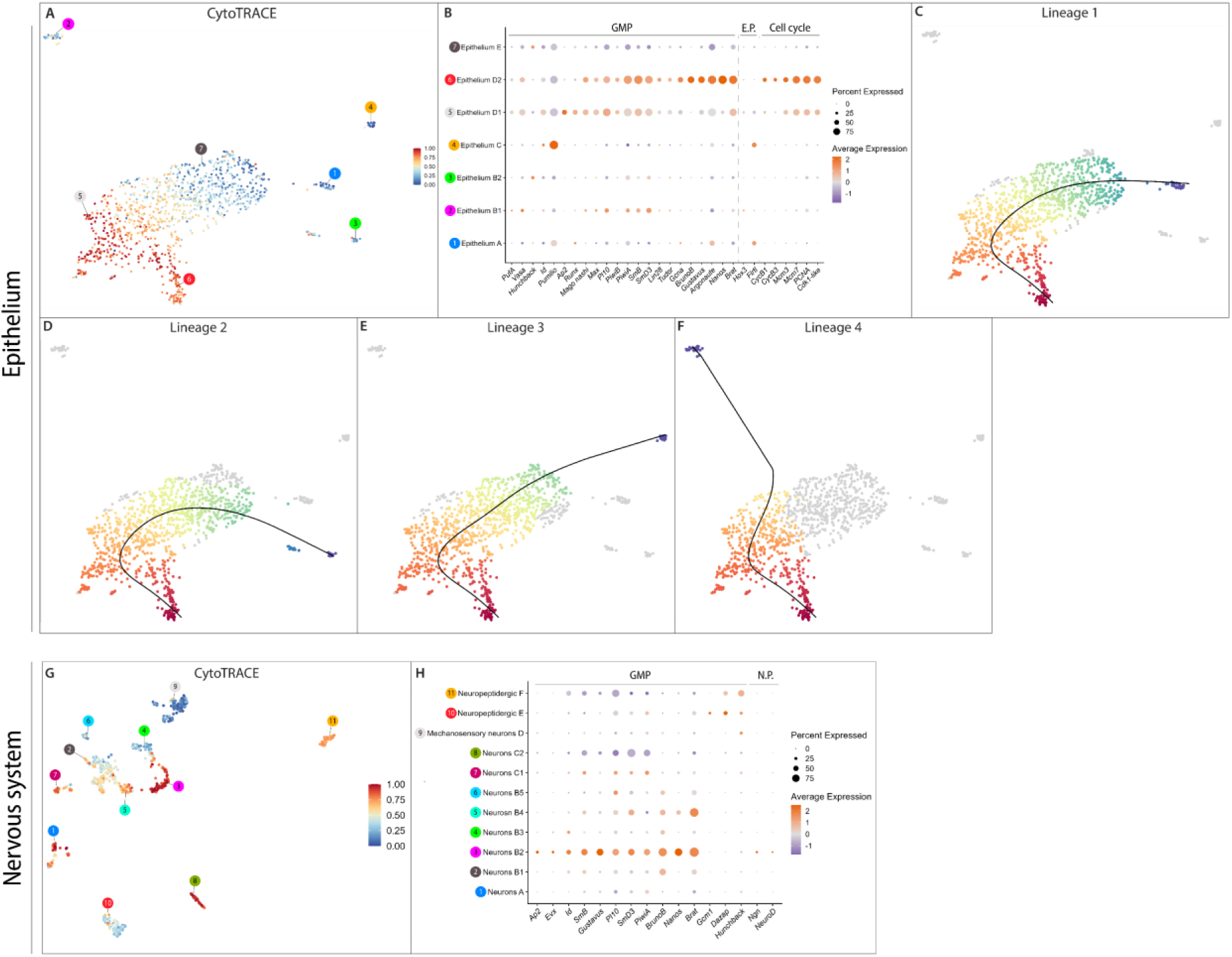
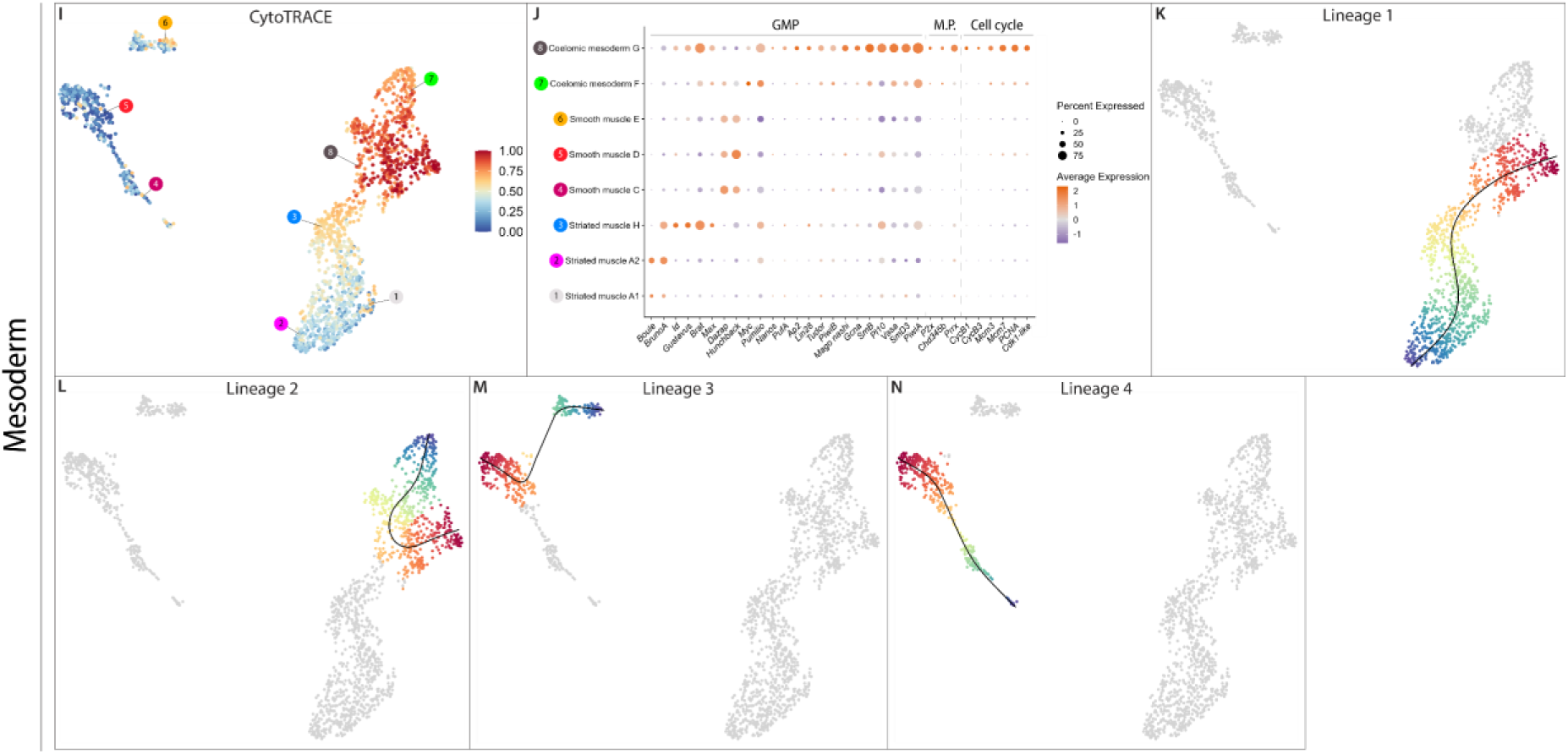
Identification of putative progenitors and their differentiation trajectories in the parapodial blastema. (A) UMAP visualization of CytoTRACE values calculated for the epithelial populations of the blastema. High values (red) indicate less differentiated cell types. (B) Dotplot of overexpressed genes belonging to the GMP signature, ectodermal progenitors, and cell cycle genes per cluster. (C-F) UMAP visualization of the Slingshot pseudo-temporal ordering of epithelial cells. Cells are colored based on their progression along pseudo-temporal space from progenitors (red) to the end of differentiation (violet). Cells outside of a given trajectory are grey. (G) UMAP visualization of CytoTRACE values calculated for the neural populations of the blastema (H) Dotplot of overexpressed genes belonging to the GMP signature and markers of neural progenitors. (I) UMAP visualization of CytoTRACE values calculated for the mesodermal populations of the blastema (J) Dotplot of overexpressed genes belonging to the GMP signature, mesodermal progenitors, and cell cycle genes per cluster. (K-N) UMAP visualization of the Slingshot pseudo-temporal ordering of mesodermal cells. E.P., ectodermal progenitors; M.P., mesodermal progenitors; N.P., neural progenitors.

**Figure S10.**
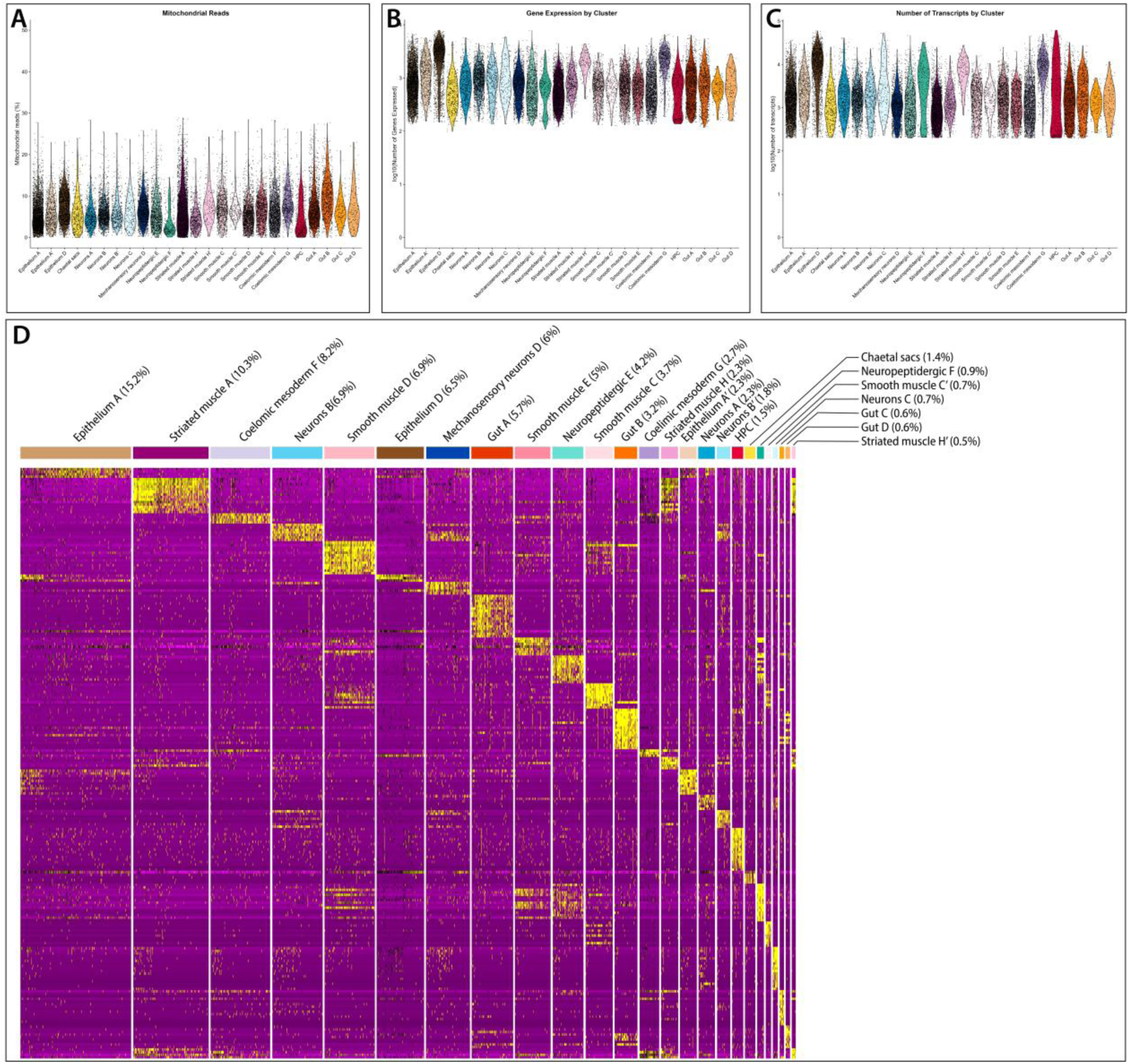
Posterior part blastema scRNA-seq quality control metrics and gene expression across different cell populations. Percentage of mitochondrial reads (A), number of genes (B), and transcripts (C) across the identified cell populations. (D) Heatmap depicting the top 20 genes expressed per cell population.

**Figure S11:**
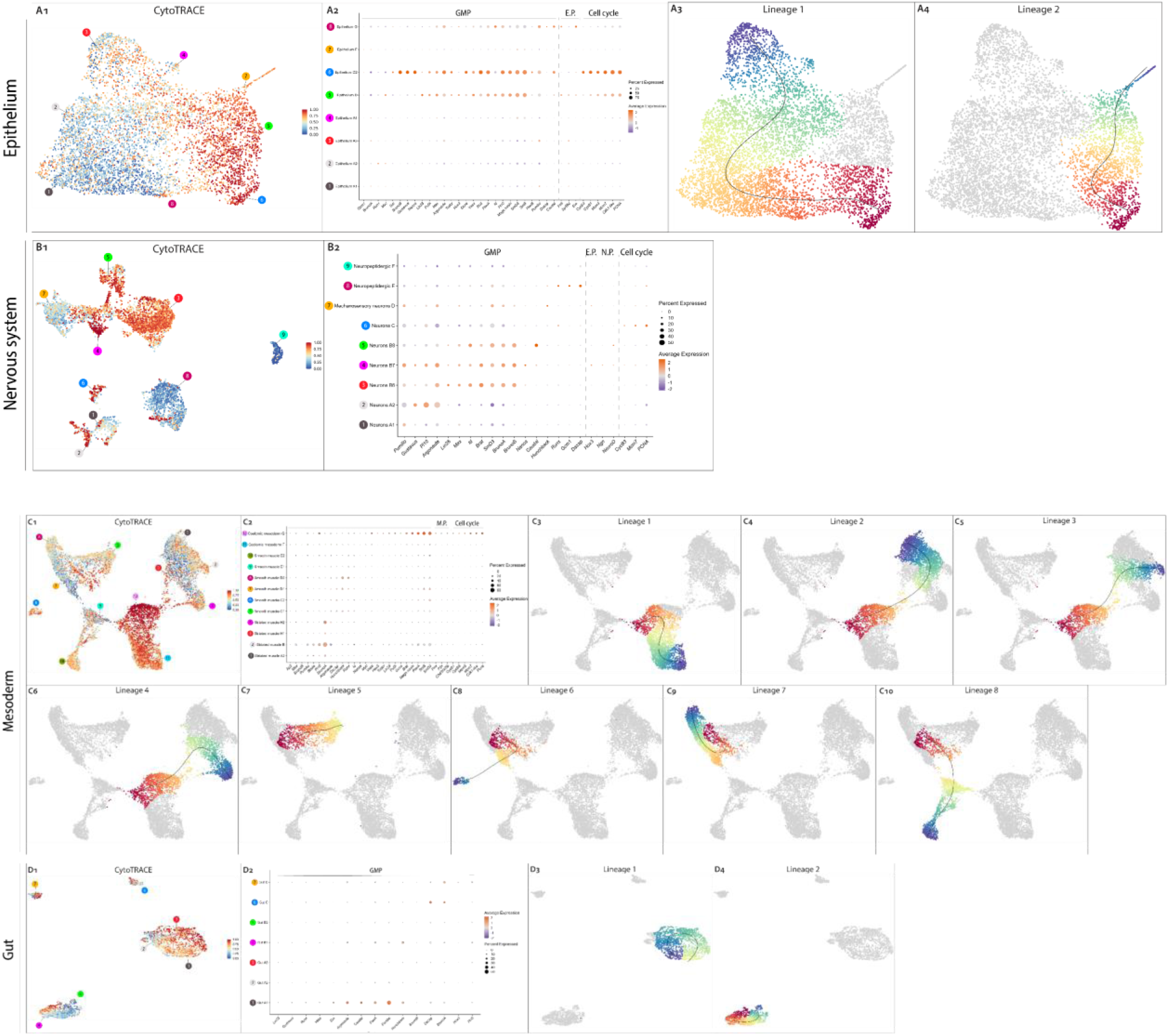
Identification of putative progenitors and their differentiation trajectories in the posterior blastema. (A1) UMAP visualization of CytoTRACE values calculated for the epidermal populations of the blastema. High values (red) indicate less differentiated cell types. (A2) Dotplot of overexpressed genes belonging to the GMP signature, ectodermal progenitors, and cell cycle genes per cluster. (A3,4) UMAP visualization of the Slingshot pseudo-temporal ordering of epidermal cells. Cells are colored based on their progression along pseudo-temporal space from progenitors (red) to the end of differentiation (violet). Cells outside of a given trajectory are grey. (B1) UMAP visualization of CytoTRACE values calculated for the neural populations of the blastema (B2) Dotplot of overexpressed genes belonging to the GMP signature, ectodermal and neural progenitors, and cell cycle genes. (C1) UMAP visualization of CytoTRACE values calculated for the mesodermal populations of the blastema (C2) Dotplot of overexpressed genes belonging to the GMP signature, mesodermal progenitors, and cell cycle genes per cluster. (C3-6) UMAP visualization of the Slingshot pseudo-temporal ordering of mesodermal cells. (D1) UMAP visualization of CytoTRACE values calculated for the gut populations of the blastema. High values (red) indicate less differentiated cell types. (D2) Dotplot of overexpressed genes belonging to the GMP signature per cluster. (D3,4) UMAP visualization of the Slingshot pseudo-temporal ordering of gut cells.

## Supplementary tables

**Supplementary Table 1.** Information on known marker genes used for cluster annotation and analysis.

| Gene | Genome annotation | Genbank | Cell type | Reference |
| --- | --- | --- | --- | --- |
| *Epig 1* | XLOC-049416 | PQ333993 | Epithelium | (Stockinger et al., 2024) |
| *Col6a6* | XLOC-015056 | PQ333995 |  | (Stockinger et al., 2024) |
| *CS3* | XLOC-001874 | KJ405470 | Chaetal sacs | (Gazave et al., 2017) |
| *Fcmg5* | XLOC_004932 | - |  | (Zakrzewski, 2011) |
| *Synaptotagmin* | XLOC-000875 | EF544397 | Neuron | (Denes et al., 2007) |
| *Rab3* | XLOC-002098 | - |  | (Achim et al., 2018) |
| *Elav* | XLOC-001549 | EF384209 |  | (Denes et al., 2007) |
| *Pkd 2-1* | XLOC-026080 | MZ647696 | Mechanosensory neurons | (Bezares-Calderón et al., 2018) |
| *Pkd1-1* | XLOC_027897 | MH035679 |  | (Bezares-Calderón et al., 2018) |
| *Qpeptin* | XLOC_000343 | KF515931 | Neuropeptigergic | (Conzelmann et al., 2013) |
| *Mef2* | XLOC-000048 | - | Striated muscle | (Brunet et al., 2016) |
| *Troponin I* | XLOC-009875 | - |  | (Brunet et al., 2016) |
| *SM-MHC* | XLOC_020862 | - | Smooth muscle | (Brunet et al., 2016) |
| *Ccdc134-like* | XLOC-046912 | PQ333994 | Coelomic mesoderm | (Stockinger et al., 2024) |
| *Egb-A2* | XLOC_024672 | MT701030 | Hemoglobin producing cell | (Song et al., 2020) |
| *Egb-A1a* | XLOC_020342 | MT701024 |  | (Song et al., 2020) |
| *Egb-A1b* | XLOC_035593 | MT701025 |  | (Song et al., 2020) |
| *Legumain protease precursor* | XLOC_037879 | KM577676 | Gut | (Williams et al., 2015) |
| *HNF4* | XLOC_039685 | - |  | (Achim et al., 2018) |

**Supplementary Table 2.** Expression data for the non-amputated parapodia dataset. DEG per cluster colored according to their over (pink) or under (yellow) expression.

**Supplementary Table 3.** Additional cell population specific gene markers.

| Gene | Genome annotation | Genbank | Cell type | Reference |
| --- | --- | --- | --- | --- |
| *Duox A* | XLOC_045030 | - | Epithelium | (Vullien et al., 2025) |
| *Acetylcholine receptor alpha9-10+* | XLOC_004519 | FJ358433 |  | (Jékely et al., 2008) |
| *Tailless* | XLOC_002678 | GU169423 |  | (Achim et al., 2018) |
| *Allatotropin receptor 1* | XLOC_037585 | KP294022 | Neurons | (Bauknecht & Jékely, 2015) |
| *Achatin receptor 1* | XLOC_023041 | KP293943 |  | (Bauknecht & Jékely, 2015) |
| *NKY-2+* | XLOC_047886 | - |  | (Conzelmann et al., 2013) |
| *Lhx1/5* | XLOC_025192 | GU169421 |  | (Tomer et al., 2010) |
| *Zic* | XLOC_064485 | - |  | (Tomer, 2009) |
| *Gata1/2/3* | XLOC_013055 | EF014968 |  | (Gillis et al., 2007) |
| *NOMPC+* | XLOC_002090 | MZ647694 |  | (Revilla-i-Domingo et al., 2021) |
| *CNGAα* | XLOC_018600 | KM199644 |  | (Tosches et al., 2014) |
| *Go-opsin 1* | XLOC_005894 | KR869765 |  | (Gühmann et al., 2015) |
| *EP* | XLOC_043838 | - |  | (Conzelmann et al., 2013) |
| *FLamide* | XLOC_056619 | - |  | (Conzelmann et al., 2013) |
| *Allatotropin* | XLOC_034858 | - | Mechanosensory neurons | (Conzelmann et al., 2013) |
| *Post1* | XLOC_037852 | DQ366678 |  | (Kulakova et al., 2007) |
| *Sll1* | XLOC_025836 | - | Neuropeptidergic | (Conzelmann et al., 2013) |
| *Uncx4* | XLOC_027840 | - |  | (Achim et al., 2018) |
| *SHM* | XLOC_034108 | - |  | (Conzelmann et al., 2013) |
| *Otx* | XLOC_060103 | AJ278856 | Nervous system related | (Vopalensky et al., 2019) |

**Supplementary Table 4.** Expression data for the non-amputated parapodia ectodermal subclustering dataset. DEG per cluster colored according to their over (pink) or under (yellow) expression.

**Supplementary Table 5.** Expression data for the non-amputated parapodia nervous system subclustering dataset. DEG per cluster colored according to their over (pink) or under (yellow) expression.

**Supplementary Table 6.** Expression data for the non-amputated parapodia mesodermal subclustering dataset. DEG per cluster colored according to their over (pink) or under (yellow) expression.

**Supplementary Table 7.** Scoring data used to establish the parapodial regeneration timeline.

**Supplementary Table 8.** EdU+ cell counting data.

**Supplementary Table 9.** Scoring data for HU cell proliferation inhibition experiment.

**Supplementary Table 10.** Expression data for the parapodia blastema dataset. DEG per cluster colored according to their over (pink) or under (yellow) expression.

**Supplementary Table 11.** Expression data for the parapodia blastema epithelial subclustering dataset. DEG per cluster colored according to their over (pink) or under (yellow) expression.

**Supplementary Table 12.** Expression data for the parapodia blastema nervous system subclustering dataset. DEG per cluster colored according to their over (pink) or under (yellow) expression.

**Supplementary Table 13.** Expression data for the parapodia blastema mesodermal subclustering dataset. DEG per cluster colored according to their over (pink) or under (yellow) expression.

**Supplementary Table 14.** New cell population specific gene markers. Genes used in the Dotplots of Figure 4 and Figure 6 are highlighted in blue.

**Supplementary Table 15.** Expression data for the posterior blastema integrated dataset. DEG per cluster colored according to their over (pink) or under (yellow) expression.

**Supplementary Table 16.** Expression data for the posterior blastema epithelial subclustering dataset. DEG per cluster colored according to their over (pink) or under (yellow) expression.

**Supplementary Table 17.** Expression data for the posterior blastema nervous system subclustering dataset. DEG per cluster colored according to their over (pink) or under (yellow) expression.

**Supplementary Table 18.** Expression data for the posterior blastema mesodermal subclustering dataset. DEG per cluster colored according to their over (pink) or under (yellow) expression.

**Supplementary Table 19.** Expression data for the posterior blastema gut subclustering dataset. DEG per cluster colored according to their over (pink) or under (yellow) expression.

## Notes

### Competing Interest Statement

The authors have declared no competing interest.

